# Why did the *Tc1*-like elements of mollusks acquired the spliceosomal introns?

**DOI:** 10.1101/656579

**Authors:** M.V. Puzakov, L.V. Puzakova, S.V. Cheresiz

## Abstract

Transposable elements are the DNA sequences capable of transpositions within the genome and, thus, exerting a considerable influence on the genome functioning and structure and providing the source of new genes. Transposable elements are classified into retrotransposons and the DNA transposons. *IS630/Tc1/mariner* superfamily of DNA transposons is one of the most diverse groups broadly represented among the eukaryotes. We identified a new group of *Tc1*-like elements in the mollusks, which we named *TLEWI*. These DNA transposons are characterized by the low copy number, the lack of terminal inverted repeats and the presence of DD36E signature and the spliceosomal introns in transposase sequence. Their prevalence among the mollusks is limited to subclass Pteriomorpha (Bivalvia). Since *TLEWI* possess the features of domesticated TE and the structure similar to the eukaryotic genes, which is not typical for the DNA transposons, we consider the hypothesis of co-optation of *TLEWI* gene by the bivalves.

## Introduction

Transposable elements (TEs) comprise a significant part of both eukaruotic and prokaryotic genomes. They represent the DNA fragments capable of transpositions within the genome and, thus, of increasing their copy number. The proportion of TEs in different genomes varies from 3% to 90% (Guo et al. 2010; de Koning et al. 2011; Sergeeva and Salina 2011). TEs play an important role in structure and function of the genomes and serve as a source of new genes (Bourque et al. 2018). A number of structural and regulatory genes has been shown to originate due to the molecular domestication of TEs (Sinzelle et al. 2009). An exposure to physical, chemical, and biological stressors can induce the TE transpositions. Genome instability due to the TE activity facilitates the origin of new features, which, in turn, can result in adaptations and future evolution (Yurchenko et al. 2011; Piacentini et al. 2014; Auvinet et al. 2018).

TEs can be classified into the retrotransposons (Class I TEs) and the DNA transposons (Class II TEs). The retrotransposons include the TEs coding for reverse transcriptase and transposing via an RNA intermediate, as well as their non-autonomous derivatives. DNA transposons represent the TEs, which do not use RNA intermediate for transposition, and their non-autonomous derivatives (Kojima 2018).

*IS630/Tc1/mariner* (*ITm*) is one of the largest DNA transposon superfamilies with its representatives in, virtually, every branch of the tree of life (Yuan and Wessler 2011). The typical size of *ITm* elements varies from one to 3 thousand base pairs (bp). Usually, those elements represent a single open reading frame (ORF) coding for the transposase of ~350 amino acid (aa) residues. The *ITm* transposases possess a domain for site-specific DNA binding in N terminal part and a catalytic DDE/D domain in C terminal part (Tellier et al. 2015). The sequences of *ITm* elements are typically flanked by a pair of terminal inverted repeats (TIRs) varying in length from 20 to 1900 bp. Some *ITm* transposons additionally have the subterminal inverted repeats (SIRs) of variable length (175-1403 bp) (Claudianos et al. 2002; Zhang et al. 2016b). Despite the variability of TIRs, their terminy usually represent a TA dinucleotide target site duplication (TSD) (van Luenen and Plasterk 1994).

*ITm* superfamily is divided into nine families: *IS630, mariner, Tc1, pogo, maT, plants, mosquito, rosa* and *L31* (Shao and Tu 2001; Tellier et al. 2015; Zhang et al. 2016b; Puzakov et al. 2018). The classification of *ITm* elements is based on phylogenetic analysis and, until recently, it was well correlated with structural features (the length of peptide chain between the second and the third amino acid residues of the catalytic DDE/D triad (Tellier et al. 2015). However, the recent studies have identified, at least, four different groups of *ITm* elements with DD37E signature: *mosquito* (Shao and Tu 2001), *TRT* (Zhang et al. 2016a), *L18* and *L31* (Puzakov et al. 2018). Thus, the families of *ITm* elements cannot be accurately distinguished by the spacer size of DDE/D domains. The study of *ITm* transposons from a Pacific oyster, *Crassostrea gigas*, revealed *Mariner-35_CGi*, which was classified as *Tc1*-like element (*TLE*) (Puzakov et al. 2018). Unlike the classic *TLEs* (DD34E), this transposon possesses a non-canonical catalytic domain (DD36E). In addition to this DD36E signature, *Mariner-35_CGi* demonstrates a number of other peculiar features, such as the lack of TIRs, an ORF split into 3-6 exons, and the presence of a full-size transcript. Our study is aiming to resolve, whether *Mariner-35_CGi* is a single copy of a particular *TLE* or it represents an unknown subfamily. An analysis of all mollusk whole-genome shotgun sequences (WGS) available at the moment of study revealed an extensive group of *Mariner-35_CGi*-like elements, which contain introns in their coding sequence. The presence of introns in Class II TEs is a rare and rather exceptional event (Laski et al. 1986; Pereira et al. 2013). Isolated cases of such introns are known among the elements of *ITm* superfamily, as the intron-bearing *ITm* transposons identified in the genome of a fungus, *Sclerotinia sclerotiorum* (Santana et al. 2014). Also, intronization was observed among the domesticated DNA transposons (Pavelitz et al. 2013; Bouallègue et al. 2017). In of our research, we describe a novel group of *TLEs*, which we referred to as *TLEWI* (***T**c1*-**l**ike **e**lements **w**ith **i**ntrons).

## Materials and methods

### Mining of DNA transposons with DD36E signature

Mining of DNA transposons with DD36E signature was performed with tBLASTn (Altschul et al. 1997). An amino acid sequence of *Mariner-35_CGi* transposase was used as a query. The nucleotide sequences of mollusk genomic DNA were obtained from NCBI WGS database (http://www.ncbi.nlm.nih.gov/). In order to identify the complete nucleotide sequences, the TE copies with highest homology to the query were derived from the respective scaffolds together with flanking areas spanning 3000 bp each. TIRs were identified in the derived sequences by BLASTn (Zhang et al. 2000). A complete sequence of each new TE with DD36E signature was used to define the element’s ends and its copy number in the genome. When determining the copy numbers, the copies with >95% query cover were counted as the full-length ones. The consensus sequences were obtained with the use of relative majority rule.

### An analysis of sequences

The identification of exon boundaries was performed visually and was guided by the maximum homology to *Mariner-35_CGi* transposase. A more accurate positioning of exon boundaries was made by a comparison with the transcribed RNA sequences of a particular TE, when available. The presence of nuclear localization sequence (NLS) was determined by ScanProsite (De Castro et al. 2006). PAIRED DNA-binding motif was identified using PSIPRED v3.3 (Buchan et al. 2013), while GRPR-type motif, G-rich box and DDE/D domain were identified visually.

### Phylogenetic analysis

Transposase sequences of 54 TEs representing different groups of *ITm* transposons were used for the phylogenetic analysis of newly identified TEs (Online Resource 1). Multiple alignment of amino acid sequences was performed by MUSCLE using the default settings (Edgar 2004). A search for the best model for phylogenetic analysis, and the analysis itself were performed with the use of MEGA7.0 software (Kumar et al. 2016).

## Results

### Distribution of *TLEWI* elements among the Mollusca species

In order to resolve, whether *Mariner-35_CGi* represents a single mutated version of *TLE* or a novel, previously unknown, group of *ITm* DNA transposons, we performed the search of Whole-Genome Shotgun (WGS) sequence database of 18 mollusk species. We used the amino acid sequence of *Mariner-35_CGi* transposase as a query. As a result, we identified 23 novel elements with DD36E signature and an ORF split in exons. We refer to these DNA transposons as to *TLEWI* (***T**c1*-**l**ike **e**lements **w**ith **i**ntrons), while *Mariner-35_CGi* was subsequently renamed into *TLEWI-1_CGi*. The names, features, copy numbers and reference localizations of the representatives of these new TEs are listed in Online Resource 2.

Despite a rather high number of unique *TLEWI* transposons, they were discovered in only 7 out of 18 mollusk species. These seven species represented three out of five orders of subclass Pteriomorphia (Bivalvia) - Mytilida, Ostreida, and Pectinida (Fig. 1). The WGS sequences of orders *Arcida* and *Limida* are not represented in NCBI. The genomes of three species belonging to subclass Heterodonta, which is also a part of the bivalves, turned out to be free of *TLEWI* elements. The TEs of this group are also lacking from the genomes of eight species belonging to classes Gastropoda and Cephalopoda.

**Fig. 1.**
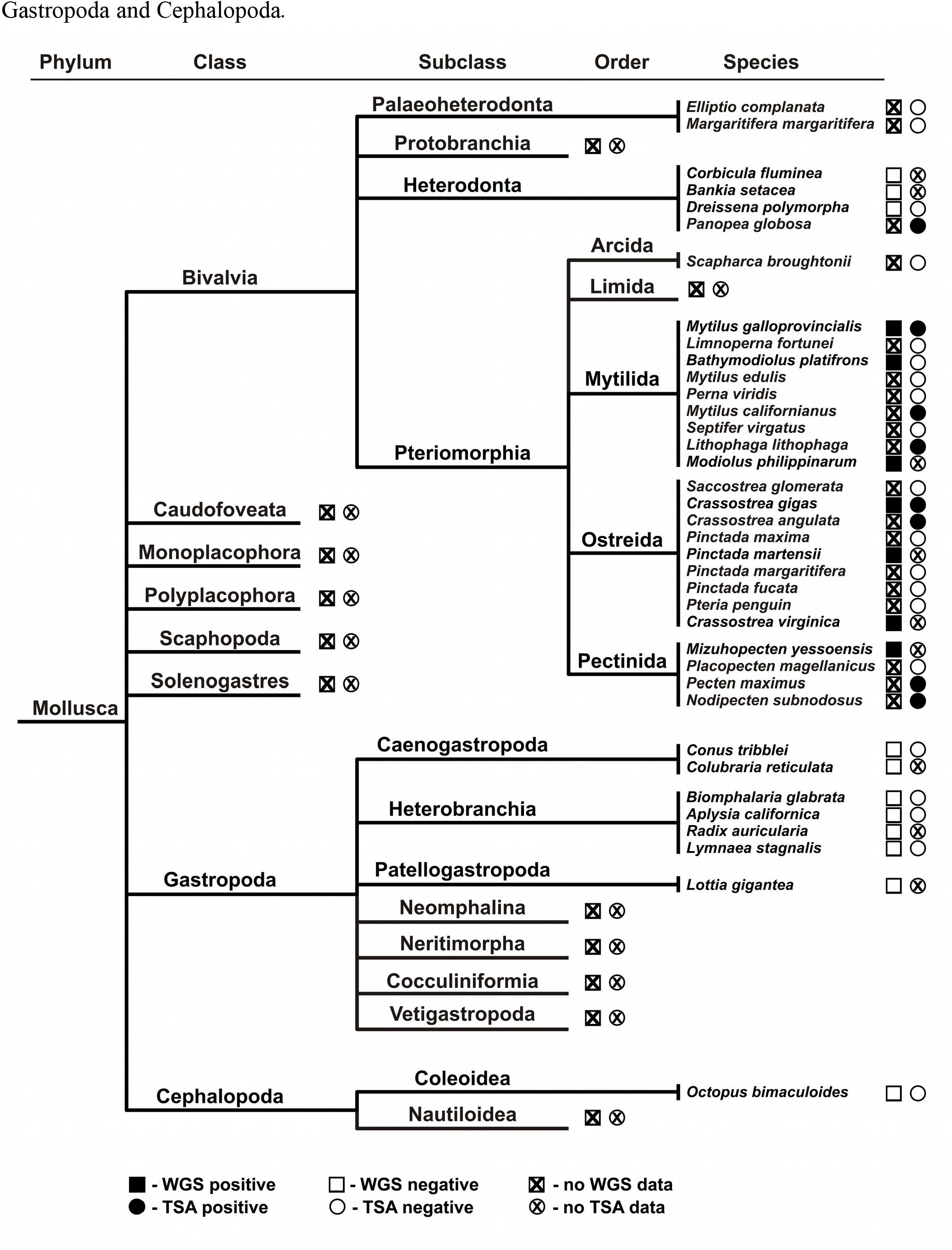
Distribution of *TLEWI* elements among the Mollusca species. The taxonomic tree of the currently living mollusk species was constructed according to the data from the World Register of Marine Species (WoRMS Editorial Board 2018)

Despite the limited distribution of *TLEWI* elements among the mollusks, the former show a considerable intraspecific variability. As much as five unique elements was identified in three species (*C. gigas, Mizuhopecten yessoensis* and *Pinctada martensii*), while three such elements were found in another two species, *Crassostrea virginica* and *Modiolus philippinarum*, and single elements were found in *Bathymodiolus platifrons* and *Mytilus galloprovincialis* (Online Resource 2). Each element was found in a varying (from 1 to 15) number of copies, with the exception of *TLEWI-1_CVi* represented by 249 homologous sequences in the genome of *C. virginica*. The number of full-size copies was not exceeding two for neither of elements. Low copy numbers of *TLEWI* elements can be due to a low transposition activity of an ancestral *TLEWI* at early stages of the transposon’s life cycle. To further investigate the distribution of *TLEWI* elements, the nucleotide sequences of the transposases were used as a BLASTn query against the species Transcriptome Shotgun Assembly (TSA) database. The transcripts homologous to *TLEWI* transposases were identified in seven species of subclass Pteriomorphia (Fig. 1). In only two of them (*M. galloprovincialis* and *C. gigas*), the new TEs were discovered by WGS analysis, while for five other species, the WGS data were lacking at the moment of study. Thus, five more species carrying *TLEWI* transposons in their genomes have been identified among Pteriomorphia. In addition, a fragment homologous to *TLEWI* transposases was identified in the mollusk *Panopea globosa* belonging to another subclass of the bivalves, Heterodonta, although *TLEWI* elements have not been discovered in the genomes of three other representatives of this taxon. We, thus, suggest that *TLEWI* distribution is not limited to subclass Pteriomorphia only or, otherwise, the above case may represent a horizontal gene transfer. Sequences homologous to *TLEWI* transposases have never been observed in the TSA databases of classes Gastropoda and Cephalopoda.

In most cases, *TLEWI*-related transcribed RNA sequences were represented by the truncated fragments mainly homologous to C terminus of the transposase. However, the transcripts spanning >90% of the full-length transposase sequence were found in four species, *C. gigas, Crassostrea angulata, Nodipecten subnodosus* and *Pecten maximus* (Online Resource 2). The transcript derived from *N. subnodosus* carried a frameshift ORF, while lacking the start codon, and, therefore, was, probably, non-functional, however, we were able to reconstruct the transposase of *TLEWI-1_NSu* element on the basis of this sequence. The transcript encoded by an ORF of a putative element of *C. angulata, TLEWI-1_Can*, was also lacking stop codon. The transcribed RNA sequence from *P. maximus* is potentially functional, which allowed us to reconstruct a complete amino acid sequence of the *TLEWI-1_PMax* transposase. Full-size transcripts of only 4 *TLEWI* out of five such elements present in the *C. gigas* have been identified. Similarly to the sequence of *TLEWI-5_CGi* element, its transcript carries the duplication of exon 3, which, probably, precludes from the translation of a functional protein. Thus, only four out of 29 TEs analyzed in this work have the potentially functional transcripts.

### The features of *TLEWI* elements

A characteristic feature of *TLEWI* elements is the lack of TIRs and TSDs, with the exceptions of *TLEWI-1_MGall*, for which the hypothetic 35 bp inverted repeats were discovered, and *TLEWI-1_CGi*, for which the 5’ TIR has only been identified based on the homology to a truncated copy of *Mariner-N1_CGi*. For this reason, the ends of *TLEWI* elements were defined as the consensus ends of their copies. For six *TLEWI* elements, the nucleotide sequences coding for the N-terminus of the transposase have not been identified, thus, precluding from the establishment of the full-length copy size. The transposons, for which the ORF ends and the expected size of full-length copies are known, are greatly varying in length, ranging from 2160 bp to 11460 bp (Fig. 2). The ORFs coding for the potentially functional transposases have been discovered for nine elements only, while the coding sequences of the latter carried either truncations, duplications, frameshifts or stop codons. The transposases of 4 elements have been reconstructed on the basis of merged sequences of two available copies (for *TLEWI-4_CGi* and *TLEWI-1_BPl*) or by the removal of duplicated loci, as in case of *TLEWI-5_CGi* or *TLEWI-3_MPh*. The transposase ORF size varies from 322 to 378 a.a. (Fig. 2), while the number of exons ranges from 3 to 6 (Fig. 3). Although the coding sequences of different *TLEWI* transposons are splitted into exons in a conservative manner, they show some differences, as well, and we, thus, identified 4 presumptive regions (A, B, C, D) (Fig. 3) in the transposase sequence for the convenience of description. The most conservative loci are C and D, which correspond to the last two exons. Exceptions to that are represented by *TLEWI-5_CGi* element, for which C locus is making a single exon with the part of locus A and the locus B, as well as *TLEWI-3_CGi*, which contains an extra exon downstream to the exon corresponding to locus D in its ORF. In eight elements, the loci A and B are merged into a single exon, while in *TLEWI-4_PMa и TLEWI-4_MYe*, A locus is split into 3 or 2 exons, respectively. Thus, the size variability of *TLEWI* transposons is mainly due to the differences in the non-coding regions.

**Fig. 2.**
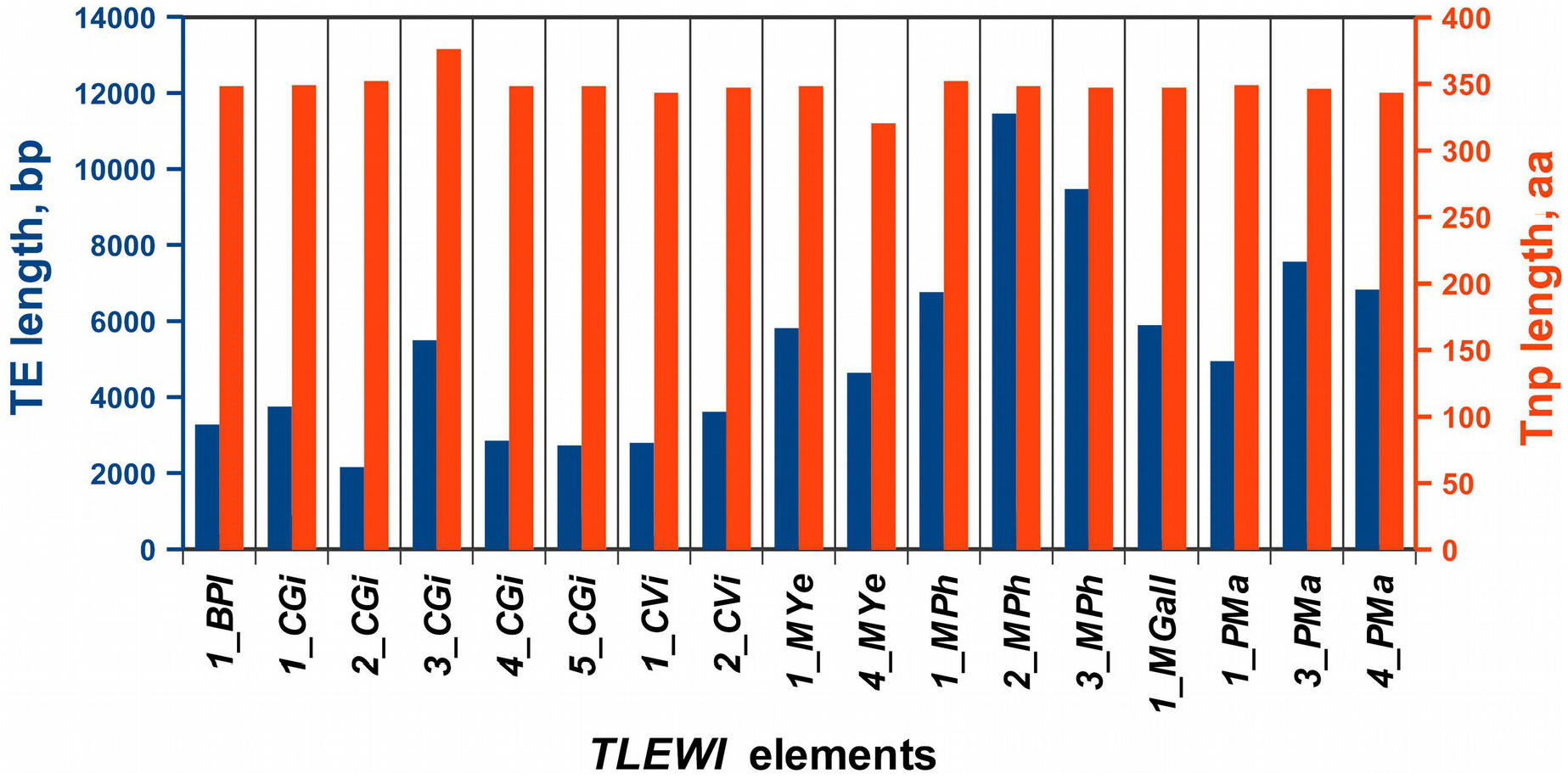
Sizes of full-lenth *TLEWI* elements and their transposases

The *TLEs* transposases typically possess two helix-turn-helix motifs (PAIRED domain), GRPR-like motif (known as an AT-hook), nuclear localization signal (NLS) and DDE domain with glycine-rich box (G-rich box) (Ivics et al. 1997; Plasterk et al. 1999). *In silico* prediction with PSIPRED helped us identify six α-helices of PAIRED domain at the N-termini of the full-length *TLEWI* transposases. Interestingly, the localization of RED sub-domain in *TLEWI* transposases is more conservative, than that of PAI subdomain, for which the boundaries of three α-helices are “blurred” (Online Resource 3). In *TLEWI-4_MYe*, the third α-helix of PAI sub-domain was completely non-identifiable, most possibly, due to a deletion between PAI and RED sub-domains. The GRPR-like motif, which is typically present in *TLEs* and is, presumably, responsible for the PAIRED domain binding to minor DNA groove at TA dinucleotide, could not have been found in neither of *TLEWI* transposases. Usually, GRPR-like motif is located between PAI and RED subdomains (Ivics and Izsvák 2015). The presumptive nuclear localization signal (NLS) was predicted (although with low confidence) by ScanProsite in only three elements (*TLEWI-1_MGall, TLEWI-1_MPh*, and *TLEWI-1_MYe*). In *TLEs* transposases, NLS is usually localized in a loop between PAIRED and DDE domains and is functional. Also, NLSs were found *in silico* in several *mariner-like* elements (*MLEs*) transposases, however, their functionality have not been corroborated (Brillet et al. 2007).

**Fig. 3.**
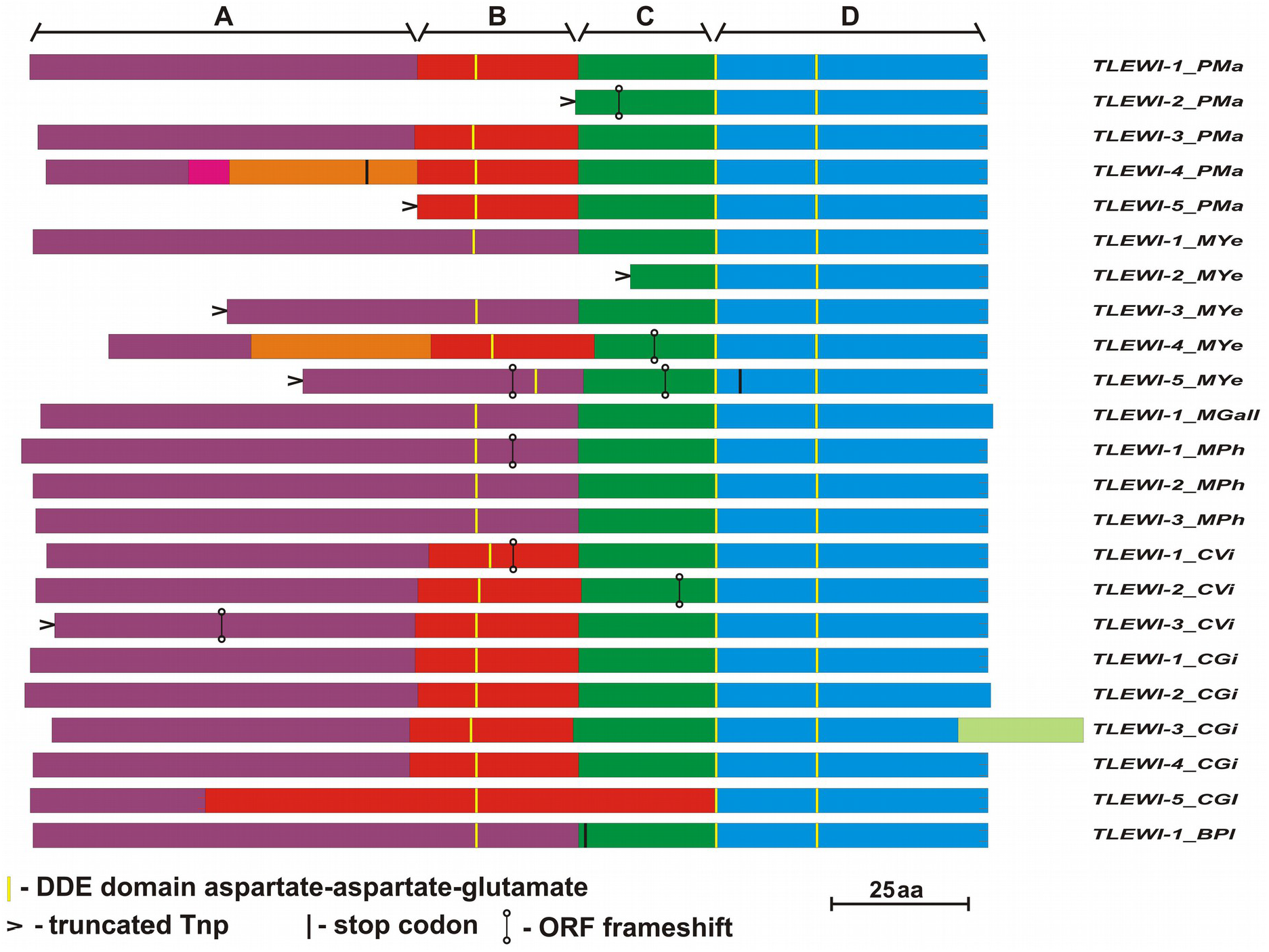
The putative *TLEWI* transposases. Four presumptive regions (A, B, C, and D) are designated in the amino acid sequences. The regions encoded by different exons are shown in different colors

The catalytic domain with DD36E signature was identified in C-terminal parts of all *TLEWI* transposases (Online Resource 3). The vast majority of elements harbor a recognizable YQDNDSKH motif near the second aspartate residue, except for *TLEWI-5_CGi, TLEWI-4_MYe* and *TLEWI-1_PMax*, in which arginine (R) substitutes for serine (S). The span between the first and second aspartate residues varies from 81 to 89 a.a., but most often it has a length of 87 a.a. Besides, in this locus we identified the G-rich box, which is a feature of *TLEs* transposases (Ivics et al. 1997). Compared to other *TLEs* transposases, the *TLEWI* G-rich box is much poorer in glycine residues. G-rich box have never been found in *MLEs* transposases or retroviral integrases, while it is an important part of functional *TLEs* transposases. Nevertheless, the function of G-rich box still remains to be determined (Ivics and Izsvák 2015). Therefore, it is currently difficult to predict, whether the reduced number of glycine residues can compromise the function of *TLEWI* transposases.

The results of multiple alingnment of full-size transposase sequences indicate that C-terminal sequences (including DDE domain) are more conserved, than N-terminal regions harboring PAIRED domain (Online Resource 3). This may indicate that DDE domain is less robust to amino acid substitutions, while sequence coding for PAIRED DNA-binding domain is more variable. The fact that N-terminal part of the transposase is not conserved within the *ITm* superfamily has been reported previously (Brillet et al. 2007).

### Evolutionary relationship between *TLEWI* elements and *ITm* transposons

In order to determine the evolutionary relationship between *TLEWI* elements and *ITm* transposons with DDE signature, we performed phylogenetic analysis, which included 24 *TLEWI* transposases (with full DDE domain) and 54 transposases representing different groups within *ITm* superfamily (Online Resource 1). As a result, we established that *TLEWI* elements form a separate branch belonging to the same clade, as *TLEs* (Fig. 4). Thus, we corroborated our previous results (Puzakov et al. 2018) indicating that *TLEWI* elements represented a subfamily of *Tc1* transposons. Additionally, we compared the exon-intron organization and the length of 21 *TLEWI* elements with their phylogeny. We did not find firm correlation between the clades of *TLEWI* elements and the features of their exon-intron structure. Since the transposases of *TLEWI-1_PMax, TLEWI-1_CAn* and *TLEWI-1_NSu* elements have been identified on the basis of TSA data, the structures of their ORFs remains unknown.

**Fig. 4.**
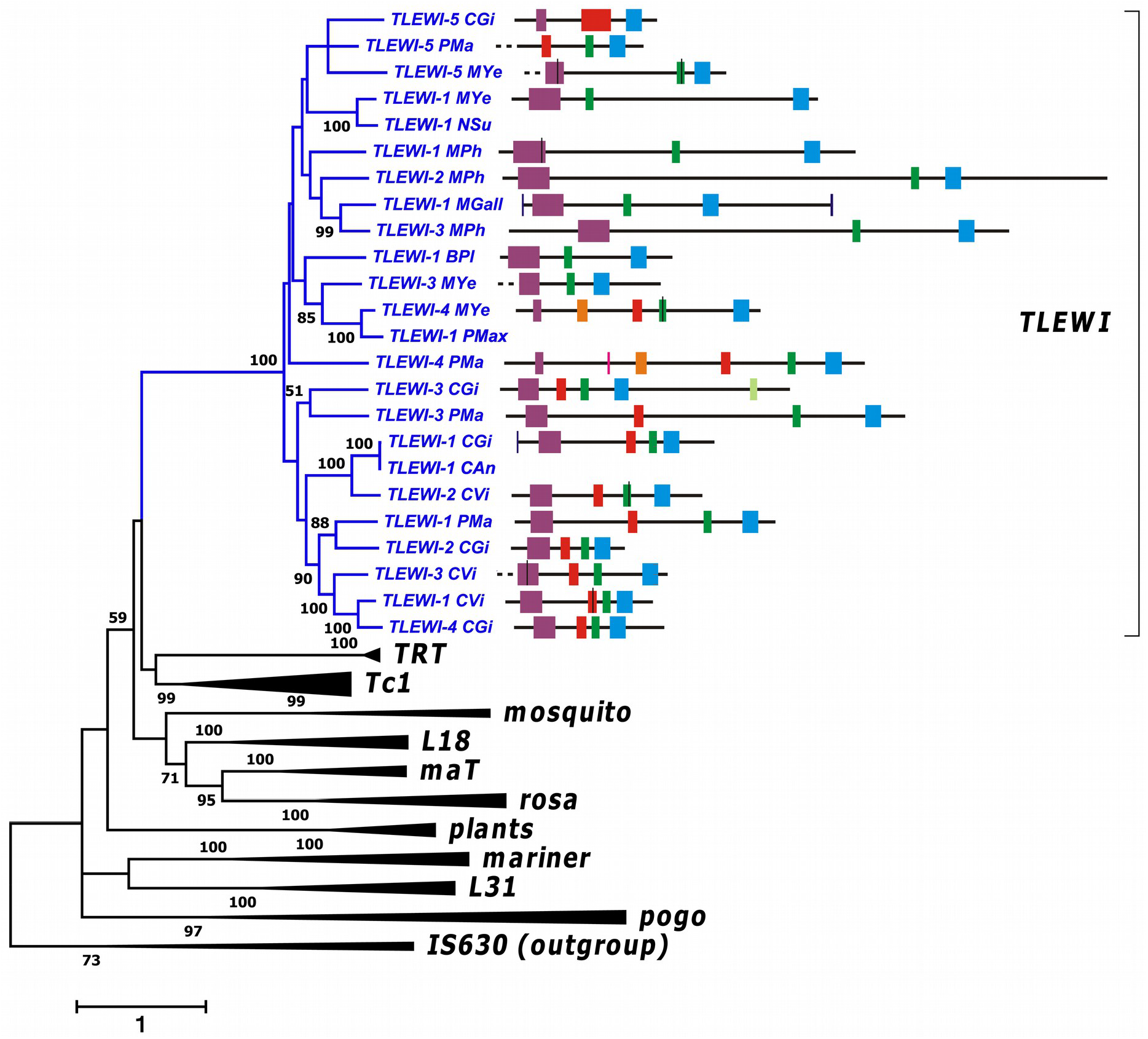
The phylogeny based on amino-acid sequences of the *IS630/Tc1/mariner* transposases. This tree was generated in MEGA7 with the Maximum Likelihood method, using the LG+G+I+F model. Only bootstrapping values higher than 50% are written on the branch. The analysis involved 78 amino acid sequences (Online Resource 1). Families are indicated in the right-hand part of the tree. The exon-intron structure of the encoding nucleotide sequences of *TLEWI* transposases is shown

## Discussion

### Distribution of *TLEWI* elements within the taxa of mollusks

The present study identifies a new group of *TLEs* in mollusks, hereinafter referred to as *TLEWI*. The specific features of these elements are their DD36E signature and an ORF segmented into exons. An analysis of genomes of 18 species belonging to the three classes of mollusks (Bivalvia, Gastropoda, Cephalopoda) revealed the new TEs in an only subclass of the bivalves, Pteriomorphia (7 species). An additional analysis of TSA databases helped us identify the presence of full-size transcripts or their fragments in another six species. Five of the latter also belong to Pteriomorphia, while the sixth, *Panopea globosa*, represents another subclass, Heterodonta. We, thus, established that the distribution of *TLEWI* elements is not limited to a single subclass.

According to the recent phylogenetic studies of the bivalves, subclass Heterodonta includes three clades (Archiheterodonta, Anomalodesmata and Imparidenta) and represents an evolutionary branch, which originated later, than Pteriomorphia (González et al. 2015). The WGS data for two subclasses, Palaeheterodonta and Protobranchia, as well as the TSA data for the latter, are not available, as yet (Fig. 5). The lack of TLEWI-related transcribed RNA sequences (as in case of Palaeheterodonta) cannot firmly testify to the absence of these TEs in the genome. If the emergence of the “first” *TLEWI* transposon corresponded to the origination of Bivalvia or Autobranchia, the TEs or their fragments would be identifiable in the respective subclasses. However, the study of three genomes of Heterodonta species did not reveal *TLEWI* elements, while in subclass Pteriomorphia the new TEs were discovered in all genomes studied. We, thus, suggest that *TLEWI* lineage have emerged in the genome of an ancient ancestor of Pteriomorphia, while the occurrence of *TLEWI* in the genome of *P. globosa* may be due to the horizontal transfer. The latter represents a rather common feature of *ITm* superfamily elements (Dupeyron et al. 2014; Wallau et al. 2016).

**Fig. 5.**
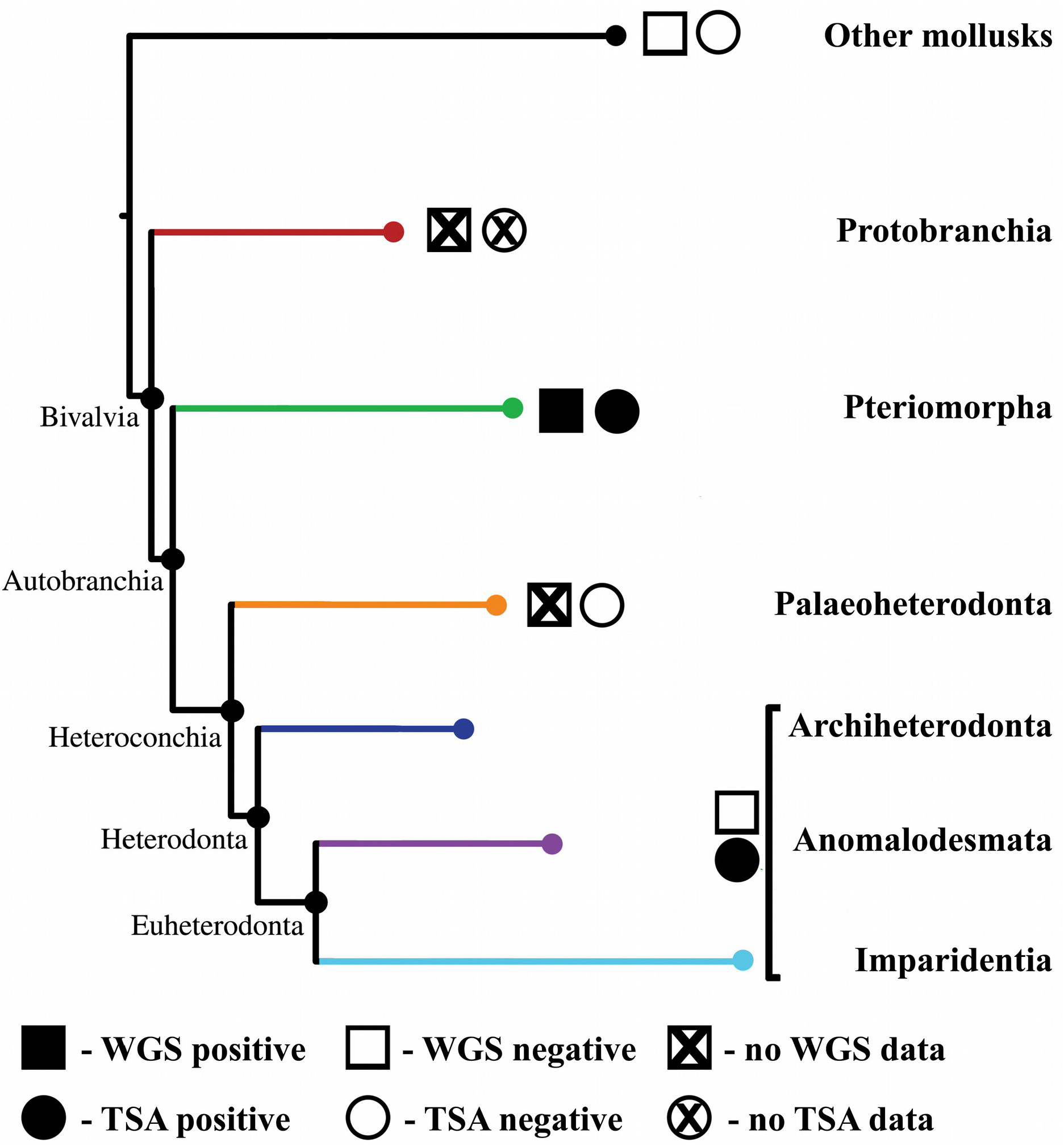
The phylogenetic tree of the bivalves by Gonzales et al. 2015 with amendments and supplements

### Intronization of *TLEWI* transposons

Currently, four types of intragenic non-coding sequences are known, such as group I and group II self-splicing introns, spliceosomal introns and tRNA introns. The types of introns are classified according to their structural features and phylogenetic distribution (Irimia and Roy 2014). Regardless of their varying lengths, virtually, all intronic sequences of *TLEWI* transposons are flanked by 5’ GT and 3’ AG dinucleotides. The structure of most spliceosomal introns is also characterized by a GT dinucleotide 5’ splice site boundary and a AG dinucleotide 3’ splice site boundary (Irimia and Roy 2014). Spliceosomal introns are separating the exons in eukaryotic genes and are excised from the RNA precursors by splicing. The finding of TLEWI-related transcribed RNA sequences without intronic regions confirms the excision of these fragments from *TLEWI* transposase pre-mRNA in the course of maturation. We, thus, classified *TLEWI* introns as the spliseosomal ones.

An analysis of gene orthologs among the animals, plants, and protests revealed numerous shared intron positions suggesting that many introns have been conserved since the time of the last eukaryotic common ancestor (LECA). A subsequent evolution involved, mainly, the loss of introns, while only several episodes of increased number of introns are known. An exon-intron structure of the coding genes was, obviously, evolving together with a eukaryotic cell, and the introns represented a major evolutionary factor throughout the entire eukaryotic history (Rogozin et al. 2012).

*TLEWI* distribution is limited to subclass Pteriomorphia only. The organisms of this subclass belonging to the bivalves, have emerged significantly later, than LECA and, therefore, the intronization of *pre-TLEWI* transposons is not linked to LECA intronization.

Spliseosomal introns are proposed to originate from Group II introns inherited from an endosymbyotic bacterium, which was the mitochondrial ancestor. The mechanism responsible for the wide distribution of spliseosomal introns in eukaryotic genomes still remain unclear. However, the dynamic processes of invasion and proliferation of spliceosomal introns were shown to resemble the life cycles of mobile introns (Group I and II introns) (Wu et al. 2017). In fungi, the emergence of new spliceosomal introns was demonstrated to result from the transposition of intron-like elements (Collemare et al. 2015). Recently, the transpositions of introns into the yeast genes *RPL8B* and *ADH2*, which preserved the activity of these genes, have been demonstrated experimentally (Lee and Stevens 2016).

The amount of intronic sequences in different *TLEWI* is varying from 2 to 5. Two scenarios are possible, due to which such variability could have emerged. The first one proposes a rapid intronization of an ancestor *TLEWI* transposon and a subsequent loss of some introns in some of them in the course of divergence of the bivalves. The second one suggests a progressive insertion of introns in the process of evolution. Based on structural (Fig. 3) and phylogenetic (Fig. 4) analysis, we consider the introns between A and B, B and C, and C and D loci of transposases to be the most ancient, since the ORF breaks in these regions are rather conservative. The introns that separate the coding sequence of some *TLEWI* in locus A have, apparently, emerged later. Thus, it looks likely that both scenarios have been involved in the generation of variability of *TLEWI* exon-intron structure.

An analysis of *TLEWI* copies demonstrated that these elements retained transposition activity after the intronization, as suggested by the identical exon-intron structure of coding sequences of different *TLEWI* copies. Intronization is important for the TEs, since their ability to transpose becomes dependent on a host splicing machinery to some extent (Laski et al. 1986, Rebollo et al. 2012). Such a dependence can become a prerequisite for the cooptation of intron-bearing TEs by the host genome. Intron containing genes are known among those originated due to the domestication of *piggyBac* superfamily elements. For example, 6 to 7 introns are found in *PGBD5* gene (in different organisms). Another two co-opted genes, *PGM* and *TPB2* harbor 2 and 12 introns, respectively (Pavelitz et al. 2013; Bouallegue et al. 2017).

### Molecular domestication and *TLEWI* transposons

The term “molecular domestication” refers to a process of co-optation of TE nucleotide sequences, which start performing some function valuable for the host genome. Molecular domestication is suggested to be the “maximum task” of TE and genome co-evolution (Bowen and Jordan 2007; Jangam et al. 2017). As a result, the TE sequences become indispensable for the host genome, while the TE itself becomes “evolutionary immortal”, as opposed to its degradation and elimination, typically, completing their life cycle.

Currently, a number of identified genes are known to be the TE “descendants”, however, for many of them, their biological function remains unclear (Sinzelle et al. 2009). Some TE proteins have been co-opted into the host defense systems protecting its genome from infectious or invasive agents (viruses and the TEs themselves) (Jangam et al. 2017). A prime example of TE-derived genes is provided by *Rag1* and *Rag2* genes. Their products Rag1 and Rag2 are the key components of specific immune response, which catalyze V(D)J recombination (Kapitonov and Jurka 2005; Panchin, Moroz 2008). The cases of domestication are known among the *ITm* transposons, as well. For example, in primates, SETMAR gene is coding for a chimeric protein, which possesses the N-terminal domain of histone-lysine N-methyl transferase and the C-terminal domain of *mariner* transposase. SETMAR protein is involved in the DNA repair mechanisms including the non-homologous end joining and the double strand break repair (Kim et al. 2014). In addition, three cases of convergent domestication of *pogo*-like transposases resulting in the emergence of centromere-binding factors have been revealed in multicellular organisms (Mateo and González 2014).

Those TE, that have undergone domestication, usually, lose the features of DNA transposons, such as the inverted repeats or the target site duplication. Moreover, the co-opted genes of these TE are, typically, present in the genomes as single copies, while their orthologs are found in other species (Feschotte and Pritham 2007). The lack of paired TIRs or TSDs, virtually, in every representative of *TLEWI* transposons indicates that they have lost their capacity to transposition. Only nine out of 29 *TLEWI* transposons possess the transposase gene lacking evident lesions (stop codons, frameshifts, duplications or deletions), and only four of them (*TLEWI-1_CGi, TLEWI-2_CGi, TLEWI-3_CGi, TLEWI-1_PMax*) were shown to produce a full-size transcript, which can testify to its preserved function. The ability of *TLEWI-1_MYe*, *TLEWI-2_MPh, TLEWI-1_PMa* or *TLEWI-3_PMa* to transcribe their genes still remains in question, since the TSA data were lacking for their host organisms. Similarly, the full-size transcribed RNA of *TLEWI-1_MGall* was not detected in the TSA database of *M. galloprovincialis*.

*TLEWI-1_CGi, TLEWI-2_CGi, TLEWI-3_CGi, TLEWI-1_MYe, TLEWI-2_MPh, TLEWI-1_PMa*, and *TLEWI-4_PMa* have all the features of domesticated TEs (Online Resource 2). However, the picture is not that unambiguous, since we also observe the disrupted TEs, TEs with more then one full-length copies and even the TEs with paired TIRs (*TLEWI-1_MGall*). An ancient *TLEWI* gene was likely co-opted by the genome, but it obtained the function, which is dispensable (neutral) for the host. In the subsequent evolution, its functional alleles were conserved only in isolated representatives of Pteriomorpha, while the non-functional orthologs remain in every representative of the taxon, as pseudogens

## Conclusion

Our results provide a pioneer *in silico* evidence for the presence and diversity of a new group of *Tc1*-like elements, referred to as *TLEWI*, in the mollusks. We determined that the distribution of these elements among the mollusks is limited to subclass Pteriomorphia. We suggest that *TLEWI* elements emerged as a result of their evolution in the genome of an ancestor of subclass Pteriomorpha, however, this hypothesis can be modified by the publication of new WGS or TSA data on the bivalves. *TLEWI* are characterized by a low copy number, the lack of TIRs, and the presence of DD36E signature and the spliceosomal introns in the transposase coding sequence. Also, some representatives possess a full-size transcript, which can be an evidence of a functional transposase gene, to some extent. Intronization of *TLEWI* transposons occurred later than LECA intronization, and, thus, the genomes of the bivalves possess (or possessed) the mechanisms facilitating the emergence and amplification of spliseosomal introns. Intron insertions into the TE coding regions make their transposition activity somehow dependent on the host splicing machinery, which can be a prerequisite for the domestication of intron-bearing TEs. Since *TLEWI* possess the features of domesticated TE and the structure similar to the eukaryotic genes (which is not the case for the DNA transposons), we suggest a hypothesis of co-optation of *TLEWI* gene by the bivalves. Data analysis will lay the ground for the future research aiming at understanding the TE roles in adaptation and evolution of the organisms.

## Supporting information

Online Resource 1

Online Resource 2

Online Resource 3

## Abbreviations

bp: base pair
aa: amino acid
TE: transposable element
ORF: open reading frame
TIR: terminal inverted repeats
SIR: subterminal inverted repeats
TSD: target-site duplication
*TLE*: *Tc1*-like element
MLE: *mariner*-like element
WGS: whole-genome shotgun sequences
TSA: transcriptome shotgun assembly

## Supplementary materials

**Online Resource 1.** Amino acid sequences of ITm transposases which were used for phylogenetic analysis

**Online Resource 2.** The features of TLEWI elements. Only copies with a query cover of over 10% were taken into account when counting the number of copies. Homologies for introns only are ignored. The copies with query cover over 95% were counted as the full-length ones. Reference transcribed RNA sequence is indicated by red letters. For the *Mytilus galloprovincialis*, two WGS projects were presented: APJB and LNJA. The results for both WGS projects are shown through slash (APJB/LNJA). TIR – terminal inverted repeats. Tnp – transposase. ORF – open reading frame. bp – base pair. aa – amino acid. “ND” – no data. “-” – not detected

**Online Resource 3.** Multiple alignment of *TLEWI* transposase sequences. Three α-helices of PAI sub-domain are typed in pink and three α-helices of RED sub-domain are typed in blue. The putative NLS is shaded in gray. The DDE triad of the catalytic domain is shaded in green, and the G-rich box of the DDE domain is shaded in yellow

