## Supplementary material for "Why did the *Tc1*-like elements of mollusks acquired the spliceosomal introns?": Online Resource 1

**Online Resource 1.** Amino acid sequences of ITm transposases which were used for phylogenetic analysis

**>TLEWI-1_BPl**
MAMGKVNRLSISTRHLIIEYSKKGLSAVHIQRLLHRKYNITTTRQSIFMFVKRYSSTGVLAPATRRDNKFPRKLTDFQRRCIDMWLRHN
SELTSQALVDRLFRVFDVRVKTSYMSKVRKALGWCTRTLQYCQLISHTNKLCRLQWSLDALRSKETFDNVIFTDETSVEMGADGGAFFY
KRTSDLDFLPAKKMKPKHAYKVHV*SGISYRGRTSICIFSGIMDSVIFQNILQSNLLPFVEHQFPDGFRLYQDNDSKHVSKSTKKWMEE
HGILDKVMTTPASSPDINPIENLWSALKGHLLKEVKPKTKDELIGGIRTFWESLTKEKCCSYIDHIHRVIPFVILNGGGPSDF

**>TLEWI-1_CGI**
MAEQRGARLSVETKRRILAIRKEERSLGKIRSVLCLRHGVKVSRQGIYMFLKKWDRDQTLVRKTRVPGIGQTKLKEIHKDLIQMWQMEN
DELTCGEMREKLKDSVGLDVSPALISKVRKDLGWQAKTGNRTCQLISRKNVRERLQWCLKAVEDKEDFKNVIFTDETTVEMCSAGRLHF
FQKSSEIQKKTSRRSRPKHAYKVHVWGGISYEGRTDICMFTGTMDSVIYQGIIEKNLIPFAEKKFPNGFRIYQDNDSKHVSKSTKEWMQ
QKNLLDKIMKTPASSPDINPIENVWAAMKLYLTKTVKPKKKEELVAGIRDFWNGLSADGCQKFIDHIHKVIPAVVLNSGEQSGY

**>TLEWI-1_CVi**
MALNSRGRRMSEKIKMVIVQYLLMHLSVEAIRPKLKNFHSYVISRQGLYYFIKQWRSGKGLRRTRNNSGNSVKLKSIHLKFMDMWLAKN
NELTTEMLKRKLFEVFGINVSPSLIRKRRSELGWKTVVSKTCQLISHKNKKVRKVWCTDALKNREDFRNVVFVDESTVEMSSFHQASSL
MQKTCARRPKPKHAYKVHVWAGISYRGRTPICIFNGIMDSLIYQNILEKNFLPFVEDKFPDGYRLYQDNDSKHKSKSTQDWMAEKGMLQ
NVMTTPASSPDLNPIENVWASMKYYLQTVAKPRTKDQLIQRIRDFWGSLSTAQCGRYIDHIHKVIPVIVLNDGEASGF

**>TLEWI-1_MGall**
MALRKSSNRLSTVTRQIILKYAPKFSALKISRLLEEKYDVKTSRQSVWRFLNRFKKSKSIRDPPRRRVRGISDLHIKAIDQWLKKDNEL
TANAIADKLRETFSTNISKGYVSLIRRKLGWTAKRVKYCQLISHVNKEKRLHYAMKVLSMKEKFLNVIFVDESTVEMTSNGRIFFYKPD
SDSSQLPARKQKPKHSYKVHVWGGISYRGRTDVCIFTGIMDSVIYQQILASNFVPFVSAAFPDGYRLYQDNDSKHKSKSTKEWMRQHGV
LDSVMETPASSPDLNPIENVWAAMKQHLQNKVKPRKKDELVQGIADFWLILTPQKCSAYIDHLHRVLPYVILNHGGPSGF

**>TLEWI-1_MPh**
MRSTRSAKVFRLSIVVRNLILKYAAAYFSTVKIQKLLQTNHQVITTRHAIRKFLDRYKITGCVSDIRPKNIAKSTSKVTVFLKQMIDMW
YSQNPEQTSRMIRNRLKDLFNVDVSNRLVSRIRRELGWTTRRAKYCQLISNSNKLKRLNWCLKTLSSKDTFHNVIFVDETTVEMSSNGK
LFFYKPNSNLQRLPARKQKPKHAYKVHVWGGISYRGKTDVCCFTGVMDSITYRGILKNNFLPFVNERFPDGYRLYQDNDSKHKSKSTVA
WVTENGVIDHVMECPASSPDINAIENLWSTLKRHLGNVVKPKKKDELVNGIKEFWESLTPEDCSNYIDHIHRVLPTIVLNDGAPSGF

**>TLEWI-1_MYe**
MAALQKISRLSPELRTSIIRYSDKGYAIRKIRKTLEETNNVTVTRKTVRLCIHRYRETGSINDRQRTGAHLKITTGVRKVIQSWLSKNN
ELTSADICARLKTELNLSVSRATINRVRVDLGWVTKRLQYCQLISHKNKRVRMKWSLDALARNDTFADVIFTDETTVEMNSSGRLFFHN
TKCGEMQRVSTKKQKPKHAYKVHVWGGISYRGRTSICIFSGIMDSVIYQGILEKKLIPFAERAFPEGYRLYQDNDSKHKSASTRSWMEN
HGVLDHVLETPASSPDINCIENVWASLKYHLQTVVKPKKKDELVCGIRDFWTSLSTEQCRRYIDHIHKVIPAVILSDGGPTGF

**>TLEWI-1_PMa**
MAATKRSTRLTTRLKAKILDLRKKMSIQKVKETLEAEDNINISRQGLYAFLNANRTRSSLGRKPRGVMPNHFKLKPVHLKVMDSWINNN
PELTASKIKFKLKETFDLDISISLISLRRRHLGWTARTSARACQLISHKNKKVRMQWCLDALQSRDDFKDVIFVDESSVEMTSNGRLFF
YKPSSDIQKKCDRKPKPKHAYKVNVWAGISYRGRTSICIFHGIMDSLTYQEILKDNLLPFVEKKFPEGFRLYQDNDSKHNSRSTKAWYE
DNNLTDSIMSTAASSPDLNPIENVWASMKLYLQTVHKPKTRDELVSGIRAFWESLTPEKCAKYIDHIHKVIPYVVLNDGEASGF

**>TLEWI-2_CGi**
MAKMCKGGRLTVRLKLKILDLYERNLSVLAIKDRLQKEDSVKFSRTTIYSFLKRHKSCGILTRRRRTVNPASVKLKELHLKFIDLWMDK
NNELKASELQKRLFETFGLKVSTSLISTRRHSVLGWKSQTSNKTCQMISRKNKTVRMQWCLDVLQTRERFRDVIFVDESCVEMCANGRI
AFHQKSSDFEKKCNRIPRPKHSYKVNVWAGISYKGKTSICIFTGIMDSEIYQKILRDNFVTVTSENFPNGYRLYQDNDSKHTSKSTKAW
MEDNGILDHIMKTPASSPDLNPIENVWASMKQYLSNVHKPRTRDELVGGIKAFWASLTAAQCGRYIDHIHRVIPYIVLNKGDASGF

**>TLEWI-2_CVi**
MAASKGTRILMETKKRIILLYNEKKSAGQIQKVLLQKYGERVSRQGVYMFLKRWKEDQVLTRKKRGKSGNVKIKKVHKDLIQMWQRRND
EVTASDLQMKLLDSTGLNVSVQSIRKVRRDLGFEAKSAGRTCQLISRKNVRERLTWCLNAVDKKEDFRNVIFTDETTVEMCSTGRLFFH
QKSSDIQQKTSRRSRPKHAYKVHVWGGISYEGRTDICIFTGTMDSVTYQHILEKNLIPANKKFPNGFRLYQDNDSKHVSKSTKEWMQQR
NLLDKVMKTPASSPDINPIENVWSAMKGFLSRTVKPRKRDELVDGIKRFWEGLSAQSCQRYIDHIHRVIPIVILNDGEASGF

**>TLEWI-2_MPh**
MATSVSARLTIPSRKLIVDYKSRGYSTVKITRVLKEKHGIITTRQSVRLFLLRFKRTGSLRDNRVNQYKQNAKRTAFLLMLLNLWLSQN
NEMTSAAICHKFLTMFGMKVSTDYVCRLRKRLGWTAQRIKYCQLISHKNKHKRMEYALDCLNKSETFDNVIFVDETTVEMTNTGRLFFF
KKGSDLQKLPAKKQKPKHSYKVHVWGGISYRGRTDICVFTGIMDSIIYQGILEDCLLGFTGSVFPDGYRLYQDNDSKHKSKSTQEWMRK
ENMLDKVMEVPASSPDLNPIENVWAAMKQHLQNVVKPKKKDELVKGILDFWSSLSSDNCGSYIDHLHNVIPTVVINEGGPSGY

**>TLEWI-3_CGi**
MAPMRLDVKAQILKYREEGRGVLEICKHLRSHHGCTVSRQAIHRFLRHGSLVRRRKLVTNSARKILAIHRRFIHMWLTQNNELTACDIQ
RKLKDLFRLSVSLSCVRKVRRELGWSSNTGKYCQQISHKNKKARMEWACNAIKNKDNFRNVVFADETSVEMCSHGKLFFHQSRSGIEMK
TRKRSRPKHAYKVNVWAGISYQGKTPICIFTGIMNSVRYQQILESNLLPFVRHRGRFPGGFRLYQDNDSKHTSRSTKAWMEQAGIINSV
MKTPASSPDLNPIENVWAGMKTFLQSTAKPKTKEELVSGIRSFWRTLTARQCARYIDHVHRVIPQIRDYLTSSLSLSMAISRYPPCFPL
RHISLASLVAAMAIALVPLDQI

**>TLEWI-3_CVi**
GRRISENAKLAIIRYRKNGLSVIKIQQLLKSHHNCIVTRPGIYNFLKKWKAGGGITAKKREGMRTLLPLHLKFVDFWLSRNNELTTQVL
QRKLFEVFGIVVSCSLIRKRRRELVWTSVVSKTCQLISNKNKMVRLQWCLDALVRKEKFENVIFVDESTVEMSASGRLFFHHEGSKIEK
KCARKPKPKHAYKVHVWAGISYRGRTNICVFNGIMDSVIYQGILENNFLPFVEKKFPDGYRLFQDNDSKHVSKSTKEWMMERGILDCVM
KTPASSPDLNPIENVWASLKYYLQTVLKPRTRDQLVDGTQNFWLSLTKKQCGRYIDHIHKVMPYVVLNSGEASGF

**>TLEWI-3_MPh**
MATARTSAFRLSREARQIIIHYSSKYSALKIGRILKEKYAITTTRQSVWRFLKRYQRTRNISDIHRSAKPKRMTEFLVKTIDLWMYENT
ELTSNAIAERLRHTFDLSFSKGHISRIRRRIGWTAQRWKYCQLISHVNREKRFQYAVKMMHSKEKFENAIFVDETTVEMCSSGRLFFYK
PDSNFSQLPAKKQKPKHSYKVHVWGGVSYRGRTNICVFNGIMDSIIYQQILENNFLPFVNRAFPDGYRLYQDNDSKHTSKSTKAWMAQN
GILDNVMETPASSPDLNPIENVWAALKMHLENKVKPKKKDELVQGIVDFWTTLTPERCAPYIDHIHRVLPAVVLNHGHPTKF

**>TLEWI-3_MYe**
KVTDSHGKFIDFWIATNPNINSRELKEKFEETFGVSVSDSLMRKVRRKLGWQTRTNIQYCQLITNVNKKKRLEYSLNALGNREDFKNVI
FVDESSVEMSSSGKMYFYKNGSCFDRLPTKAPKPKHAYKVHVWAGISYQGRTNLCIFSGVMDSVIYQGILQDNLLPFVNMRFPDGYRLY
QDNDRKHTSKSTQEWMSENNMLENVMTVPPSSPDLNPIENVWAAMKDYLQRNVKPKKKEELITGIRTFWESLTPQHCARYIDHVHKVIP
HVVLNMGGPTQF

**>TLEWI-3_PMa**
MASSDRGKRLSVETRKFILARKSDGLSVQRIKSELKKCLLIDVSRQTIYNIVSHGSLVRRRSTNGGNATKIRPVHMRFLNIWLSKNNEL
TSKDIVDKLRSTFGVGISHGYARIIRRKLGWMKSTGKYCQLISNKNKAVRKAWCCDAWHRRDTFEDVIFCDESSIEMNSNGRLFFHRER
SAGFSKTTTKRSKPKHSYKVNVWAGISYRGRTPICIFTGIMDSVIFQQILQDNLLPFVQREFIDGFRLYQDNDSKHNSRSTQEWMRQVG
ILDTVMKTPASSPDLNPIENVWASLKLHLQTKVKPKRKEDLVRGIQEFWKSLTPESCRKYIEHIHKVIPHVILNNGDPTDF

**>TLEWI-4_CGi**
MVVNTRGRRMSEGIKMMIVRYLLLDLYVEVIREKLKHLHGFVISRQGLYAFKRRWQTGKELRITRKICNGAVKLKKIHLKFMDMWLAKN
NELTTGMLKRKLFEVFGVNVSTSLIRKRRSELGWNTVVSKTCKLISKKNIQVRQSWCLDALNNKEDFRNLIFVDENTVEMSFSGRLFFH
QPGSKIQKICARRPTPKHAYTVHVWAGISYRERTSICIFNGVMDSVVYQDILDKNCIPFVEERFPDGYRLYQDNDSKHKSKSTRDWMAE
KGILDNVMTTPASSPDLNPIENVWASMKYYLQTVAKPRTKDQLIQGIKAFWETLSTPQCERYIDHIHKVIPYVVLNRGEASGY

**>TLEWI-4_MYe**
MADLRRRRLSVEAKLKIMSMKSKTGGQITKILKEKYGILTTRKTVNIYRKRLVKPIHKKIINMWLSRNSELTSPEIQRKFHSSFGLTVS
TAHTRRIRRELGWTARHIKYCQLISNKNKNRYNFCINTLISLETFSNVIFVDETSVEMSSNGRMFFTQHGPSMDRLPSKAPKPKHAYKV
SYRGRTDICIFSGIMDSIIYKGILQDNLLGFVDANFRAGFRLYQDNDRKHTSRSTQDWMRKKGILDSVMSVPASSPDLNPIENVWSAMK
DFIQRQVKPKKKDELIKGIVQFWNDLTPEKCRRFIDHIHRVIPHVVLNRGGPTQF

**>TLEWI-4_PMa**
MASTGRRLTITARKLILRYKEEGLSHAKIKQVLRDRHLITTTKKSIYRILAHFELVICPLTNTYNSYKLTDLHRQVLHMWMQEKPESTS
TALSLRFKEELGINVSQSHVRLIRRQMG*KTSQAKYCQLISAPNRRVRLEWALNALSSNEKFEDVIFMDETTVEMSSTGRVFSHETVGR
LDCMANKQAKPKHAYKVHVWGGISYHGRTDICIFTGTMDSVIYQQILEKNLIPFAKRVFPNGYRLYQDNDSKHKSKSTKEWMSRNGIEK
NFLEAPASSPDINPIENLWAALKRFLSCEVKPKKKDELVEGIRTFWESLTVDTCSSYIDHIHKVLPHVVLNNGHATKY

**>TLEWI-5_CGi**
MAPGRGGRSLSLSIRNIIIKLHKAGISPVKICKTLQDKHQFKTTRQSVRRFIIRFETTGCVHDKRKPRRPEHTKVRRIHMQFINMWMAQ
NPEMTAANIQGRLEKQCGIQLSKEYVCMLRRKLNWTPKHMKYGQLISHKNVKHRLDWCLDQLISKDSFRDVIYVDESTVEMCSSGRLFF
HRHGSNMDRLPAKAPKPKHSYKVNVWGGISHRGRTSICIFSGIMDSEIYQKILEDNLLPFATANYPDGFRLYQDNDRKHTSKSTRRWME
EKQLTESVMTTPASSPDLNPIENVWSAMKTYLRSVVKPKKKEELVQGIVSFWNTLSAEKCGHYIDHVHRVIPHVVLNAGGPTQF

**>TLEWI-5_MYe**
IHLKFLEMWMEHDPDITSLSLQQKMKSILGVTVSASFICKLRKLMGWTAKHVKYCQLISHKNKQCRLDWTLNALEAKDFKNVVFVDETT
VEMCANGGLFFINLHVWGGISYHGRTDICIFTGIMDSKIYQQILENLIPWANEMFPEGLRLYDNDSKHRSR*TTEWFRSRNMEENVMAT
PASSPDLNPIENVWSALKGHLQKCVKPRTKDDLQNGIMCFWESLSARHCAKYIDHIHKVIPHVVLNKGGATVF

**>TLEWI-5_PMa**
RLSWCLNALAVKETFHDVIFVDETTVEMCSSGRLHFYHSKSEMQRLPSKQPKPKHSYKVHVWGGISYRGRTHICIFTGIMDSVIYQQIL
QRNLLTFTSSVFPEGFRLYQDNDSKHKSRSTRKWMEEVNILQNVMETPASSPDLNPIENVWSTMKYHLQTYVKPKTKDELVNGIASFWS
TLTTADCTKYINHIHRVIPHVVLNNGEPTVF

**>TLEWI-1_PMax**
MADPRSRRITMEAKLKIVSMKTKSGSEIARILKENFNILTTKKTVNVYRRRIKRTGMLEIVRSSPALKNRQLVKPLHKKIINMWLSRNS
EMTSPELQRKFHSTFGFSVSTAHLRRIRKELGWKSRHVKYCQLISNKNKINRYNFCIEALINLETFSNVIFVDETTVEMSSNGRMFFTQ
HGPSMNRLPSKASKPKHAYKVHVWGGISYSGRTDICIFSGTMDSVIYQSILQDNLLKFTATNFGDDFRLYQDNDRKHTSKSTQEWMRQN
GILDKVMVVPASSPDLNPIENVWSAMKDQIQRHVKPKKKNELISGIIDFWKGLTPEKCQRFIDHVHRVIPHVVLNRGGPTQF

**>TLEWI-1_CAn**
RLSVETKRRILAIRKEERSLGKIRSVLCLRYGVKVSRQGIYMFLKKWDRDQTLVRKTRVPGIGQTKLKEIHKDLIQMWQMENDELTCGE
MREKLKDSVGLDVSPALISKVRKDLGWQAKTGNRTCQLISRKNVRERLQWCLKAVEDKEDFKNVIFTDETTVEMCSAGRLHFFQKSSEI
QKKTSRRSRPKHAYKVHVWGGISYEGRTDICMFTGTMDSVIYQGIIEKNLIPFAEKKFPNGFRIYQDNDSKHVSKSTKEWMQQKNLLDK
IMKTPASSPDINPIENVWAAMKLYLTKTVKPKKKEELVAGIR

**>TLEWI-1_NSu**
RLSPEVRSSVVRYSIQGCGIKQIRKKLEENHHITVTRKSVRLCIQRYRETGSITNRRRRGNPKLRLTTGIRKVMQTWLSENNELTAADI
CERLKKELNVEVSMSTVNAARAELGWVAKRLQYCQLISHKNKRARVQWCLDALARKDTFTDVVFTDETTVEMNSNGRLFFHNVKCGEFQ
RTSAKRQKPKHAYKVHVWGGISYRGRTAICIFSGIMDSVIYQGILEKNLIPFAERAFPSGYRLYQDNDSKHTSASTRSWMERNGVLDHV
LKTPASSPDINCIENVWASMKYHLQSVVKPKKKDELVQGIRDFWSKLTMEQCRKYIDHIHKVLPVVILNEGGPTGY

**>DD34D Dmmar1 (X78906)**
MSSFVPNKEQTRTVLIFCFHLKKTAAESHRMLVEAFGEQVPTVKTCERWFQRFKSGDFDVDDKEHGKPPKRYEDAELQALLDEDDAQTQ
KQLAEQLEVSQQAVSNRLREMGKIQKVGRWVPHELNERQMERRKNTCEILLSRYKRKSFLHRIVTGDEKWIFFVNPKRKKSYVDPGQPA
TSTARPNRFGKKTMLCVWWDQSGVIYYELLKPGETVNTARYQQQLINLNRALQRKRPEYQKRQHKVIFLHDNAPSHTARAVRDTLETLN
WEVLPHAAYSPDLAPSDYHLFASMGHALAEQRFDSYESVKKWLDEWFAAKDDEFYWRGIHKLPERWEKCVASDGKYFE

**>DD34D Hsmar1 (U52077)**
MEMMLDKKQIRAIFLFEFKMGRKAAETTRNINNAFGPGTANERTVQWWFKKFCKGDESLEDEERSGRPSEVDNDQLRAIIEADPLTTTR
EVAEELNVDHSTVVRHLKQIGKVKKLDKWVPHELSENQKNRRFEVSSSLILRNNNEPFLDRIVTCDEKWILYDNRRRPAQWLDREEAPK
HFPKPNLHQKKVMVTVWWSAAGLIHYSFLNPGETITSEKYAQQIDEMHRKLQRLQPALVNRKGPILLHDNARPHVAQPTLQKLNELGYE
VLPHPPYSPDLSPTDYHFFKHLDNFLQGKRFHNQQDAENAFQEFVESRSTDFYATGINKLISRWQKCVDCNGSYFD

**>DD34D Famar1 (AY155492)**
MENQKEHFRHILLFYFRKGKNASQAHKKLCAVYGDEAFKERQCQNWFAKFRSGDFSLKDEKRSGRPVEVDDDLIKAIIDSDRHSTTREI
AEKLHVSHTCIENHLKQLGYVQKLDTWVPHELKETHLTQRINSCDLLKKRNENDPFLKRLITGDEKWVVYNNIKRKRSWSRPGEPAQTT
SKAGIHQKKVLLSVWWDYKGIVYFELLPPNRTINSVVYIEQLTKLNNAVEEKRAELTNRKGVVFHHDNARPHTSLVTRQKLLELGWDVL
PHPPYSPDLAPSDYFLFRSLQNSLNGKNFNNDDDVKSYLIQFFANKNQKFYERGIMMLPERWQKVIDQNGQYITE

**>DD34D Bytmar1 (CAD45367)**
MGKIEYHAVIKFLTKVGKNAKEIHDRLVAVYNDTASSYATVTRWHKEFRHGRESLEDDSRVGRTFEATSEDTVDRVEAMIMENRRVKVE
EISLEIRISHGSVCTIINHHLGMSKVSARWVPRNLSLHDRLQGQTSSEELLTLYNAYPAGFKSRVMTGDETWVHHWDPETKLESMAWKQ
KGSPTPLKFWTQPLAGKIMATIFWDAGGVLLVDVLPRGSTITGKYYAGVLGRLRDSIRQKRRGKLTRGVLLLLHDNAPVHKAHHAQAAL
RDCGFEQFNHPSYSPDLAPNDYFLFRQLKSSLRGRRFDDNDEVKEAVMMWLEEQLESFWLAGIQSPSRQVVQMYSIKGNYIEK

**>DD34D Tvmar1 (AAP45328)**
MYMMSMLYFGAMPGYKMNFETRGGFAGNFGGIFFIFLMNHKENILALAKKCKDCKKIYETLVKCFGMDAPSYSTVTYHVRMYHFMNKKA
PIIKIDKKSPDQRKIKAILQALDEDPRASLRRIEEMTKIPRTTVSYYLHNYLNYKLAYTRWVPHNLNSVQKKSRVQSSKELLSILGAYQ
SKKFRFLVTGDESWFQYATEAKIMWIPKDENPQTFPKKKIDTPMMMLSVFWGVNGIIAIDILQKPNTMNAQYLIDNVLTQIINSDEFEK
SKQQKQKFAIHFDNSRVHKSHKVMNYLVENNVKVVPNPIYSPDIAPSDFYLFGTLKKRAEGREFASPDDLENFVREQFEQFSHDDLKRV
FQAWIDRCERVIESNGDYI

**>DD34E Tc1 (X01005)**
MDRNILRSAREDPHRTATDIQMIISSPNEPVPSKRTVRRRLQQAGLHGRKPVKKPFISKKNRMARVAWAKAHLRWGRQEWAKHIWSDES
KFNLFGSDGNSWVRRPVGSRYSPKYQCPTVKHGGGSVMVWGCFTSTSMGPLRRIQSIMDRFQYENIFETTMRPWALQNVGRGFVFQQDN
DPKHTSLHVRSWFQRRHVHLLDWPSQSPDLNPIEHLWEELERRLGGIRASNADAKFNQLENAWKAIPMSVIHKLIDSMPRRCQAVIDAN
GYATKY

**>DD34E Passport (CAB51371)**
MKTKELTKQVRDKVVEKYEAGLGYKKISRALNISLSTIKSIIRKWKEYGTTANLPRGGRPPKLKSRTRRKLIREATRRPMVTLEELQRS
TAEVGESVHGTTISRLLHKSGLYGRVARRKPLLKGIHKKSRLEFARSHVGDTANMWKKVLWSDETKIELFGLNAKRYVWRKPNTAHHPE
HTIPTVKHGGGSIMLWGCFSSAGTGKLVRIEGKMDGAKYREILEENLMQSAKDLRLGRRFIFQQDNDPKHTARATKEWFGLKNVNVLKW
PSQSPDLNPIENLWQDLKIAVHRRSPSNLTELHLFCQEEWTNLSISRCAKLVETYPKRLAAVIAAKGGSTKY

**>DD34E Quetzal (AAB02109)**
MTREELSVSKRQDIIRLHGAQGKSYTEIAMLTNINRNTVARVIQRYKYEGRVSNLPRKGRPSVCTDRMRRAIKRLVDAEPEISAQSVAI
VLNERHGIAISCETVRRYIHKFGYKAYNRRKKPQISPINRKRRLEFAKKYVNHPPEFWKKVLFTDESKFNIFGWDGTIKVWRPPGEGLN
PKYTAKTVKHNGGGVLVWGCMAANGVGNLQVIDGIMDQYVYINILKQNLGPSLEKLGMSQDYWFQQDNDPKHTAFNSRLFLLYNTPHQL
KSPPQSPDLNPIEHAWELLERKIRQTRIKNRVDLENKLKEAWITISEDYTQNLVNSMPRRLAEVIKMKGYATRY

**>DD34E Frog Prince (AAP49009)**
MPRPKEIQEQLRKKVIEIYQSGKGYKAISKALGIQRTTVRAIIHKWRRHGTVVNLPRSGRPPKITPRAQRRLIQEVTKDPTTTSKELQA
SLASVKVSVHASTIRKRLGKNGLHGRVPRRKPLLSKKNIKARLNFSTTHLDDPQDFWDNILWTDETKVELFGRCVSKYIWRRRNTAFHK
KNIIPTVKYGGGSVMVWGCFAASGPGRLAVIKGTMNSAVYQEILKENVRPSVRVLKLKRTWVLQQDNDPKHTSKSTTEWLKKNKMKTLE
WPSQSPDLNPIEMLWYDLKKAVHARKPSNVTELGQFCKDEWAKIPPGRCKSLIARYRKRLVAVVAAKGGPTSY

**>DD34E Sleeping Beauty (AFR53956)**
MGKSKEISQDLRKKIVDLHKSGSSLGAISKRLKVPRSSVQTIVRKYKHHGTTQPSYRSGRRRVLSPRDERTLVRKVQINPRTTAKDLVK
MLEETGTKVSISTVKRVLYRHNLKGRSARKKPLLQNRHKKARLRFARAHGDKDRTFWRNVLWSDETKIELFGHNDHRYVWRKKGEACKP
KNTIPTVKHGGGSIMLWGCFAAGGTGALHKIDGIMRKENYVDILKQHLKTSVRKLKLGRKWVFQQDNDPKHTSKHVRKWLKDNKVKVLE
WPSQSPDLNPIENLWAELKKRVRARRPTNLTQLHQLCQEEWAKIHPTYCGKLVEGYPKRLTQVKQFKGNATKY

**>DD34E Mariner-5_CGi (Repbase, https://www.girinst.org)**
MGRKAKELSNEVKEMVWKMLQDGKTITYVSETFGIPRSTISSFKKRVEQRGYIENIPRRGRKSTVTSRDYRKLERLVKTHRRESLKDIT
DQFNENRERPVSKRTVQLHLHKNGFVRRVAKKKLVIREVNRKKRLSWCREKRKLTVGNYWRKVIFSDESKIVVGQNSRVYIWRKRGEGW
RPDLVEARRAKPSYEVMIWGCISWHGVGTITAVNGNINAEKYQQVLNDNLWPVIVRHFPKEQYIFQDDNAPVHRARSTQDFIHRNGIRT
MSWPAQSPDLNIIENLWLLLKRKLQARVGRIENKDDLFREIQRVWTSITPQYVQSLYNTIPRRLQSVIRLKGHLTKY

**>DD34E Mariner-23_CGi (Repbase, https://www.girinst.org)**
MDTKRQEMTDETKTIILELFDSGFKQSEISKVLKISPVTICRFLKRYHESGSIENRPRSGRPKKMDDRSCRTLNRIIRGNRTSTLVDIT
NEFNTFNHVSVSKRTIQRSIHKYDLRKRAIAKKLGVRQINRVRRVSWARSKLTWTVAENWSKIIFSDEMAVTVSGCGAVKVWRTKREKY
LPECIGHLKRHTGRNLKVMVWGCITYHGQGLLTFIDGTMDSEKYINVLDENLWPVIAKYFSNKPYMFQDDNAPPHASRMTKAWKEXNNI
PTLTXPAQSPDVNIIENIWLLLKNQVKRHMMNIDTVDDLKAVLQHAWLSIPLMYIQNLYSSIPRRLQLVIRSKGHITKY

**>DD34E Mariner-8_CGi (Repbase, https://www.girinst.org)**
MPPRKLEEIPVAVRMKVIEAYKTEKNKAEIGRLLGIPRTTIVSIIKKFEVHGSVENRHRSGRKPKFSVRDEVQLSRIMKSNRRGTLDDI
TSEINNAKDSTFCSKTVRRKLFKMGYNRRVQKKKMVVRVCNRKKRVSWCRERKNWTVDDHWKNWIFSDECQVVIGDNNRVYIWRKTDEV
DNPHLVCAPSKSKLSVMIWGCICYDGVGTLTSVNGNINSEKYIDILENNLWPVIVRHFPHGNYVFQDDNAPVHRSRLLSAYMEENNINT
TSWPAQSPDINIIENIWLKLKRHLQSTRNTIDSKDQLIAEVTRFWQNLSVDYIRQLYDTIPTRLXEVIKMKGNLTKY

**>DD34E Mariner-14_CGi (Repbase, https://www.girinst.org)**
MGRKGSELCASEKESILSLSKANIKLKEISEITGRPISTICSFLKRQERRGITENKNRSGRPRKCSVQGERQLIRLVKNNRRRTLTELT
NVSNENGASRLSESTVKRILRKGGYRRRLVKKKLRIREVNKKKRVNWCKQYRHKTVDDFWNNIIFSDECKVMIDGEQRVYVWRKDGEEW
DPPCVAPPPGRRLDLMVWGCITAHGVGTLCIVDGNINAEKYIEIIDSNLWPVVAMHFPNNQYIFQDDNAPVHRARVVKDFVTREGITTL
EWPAQSPDLNIIENCWKKMKHEINRNVHNLRTNDDLAAAVRQAWENIPLQFIQRLYQSIPRRIQAVIKSKGCLTKY

**>DD37D Bmmar1 (U47917)**
MEWGDKENRIAVIALHKVGMEPNAIFKTLHTLGISKMFVYRAINRCNETSSVCDRKRSGRPRSVRTKKVVKAVRERIRRNPVRKQKILS
REMKIAPRTMSRILKDDLGLAAYKRRTGHFLTDNLKENRVVKSKQLLKRYAKGGHRKILFTDENFFTIEQHFNKQNDRIYAQSSKEASQ
LVDRVQRGHYPTSVMVWWGISYEGVTEPYFCEKGIKTSAQVYQDTILEKVVKPLNNTMFNNQEWSFQQDSAPGHKARSTQSWLETNVSD
FIRAEDWPSSSPDLNPLDYDLWSVLESTACSKRHDNLESLKQSVRLAVKIFPMERVRASIDNWPQRLKDCIAANGDHFE

**>DD37D Bmmar6 (AF461149)**
MEWTLKEDRVAVIALHRCGYAPIQIFNILKNLNITKRFVYRTIKRYNEDSSVDDRSRSGRPRSVRTPAVIKAVKARIQRNPKRKQKLLA
LQMGLSRTTVKRVLNEDLGLRAYRRKTGHRLNARLMDLRLKRCRALLKRYAGKKYREILFSDEKIFTVEESYNKQNDKVYAHSSEEASN
RIPRVQRGHFPSSLMVWLGVSYWGLTEVHFCEKGVKTNAVVYQNTVLTNLVEPVSHTMFNNRHWVFQQDSAPAHRAKSTQDWLAAREID
FIRHEDWPSSSPDLNPLDYKIWQHLEEKACSKPHPNLESLKTSLIKAAADIDMDLVRAAIDDWPRRLKACIQNHGGHFE

**>DD37D Cemar6 (LK928390)**
MRASPMREPIVRFHRNGVAAKSIARRLKVSEKLVSTTIARFKELGNFSDRSGRGRPPTVTTPAMIKKVRGRFRHNSGRSVRAMARELKI
SQSSLCRMVKNNLKLKAYKKSTCQFLSEAAKIKRKDRAMNLLRRFRNGAHRKVLFTDEKIFCIEQSFNTQNDRVYAKTQPNSRVQRTGY
PKGIMVFAGITANGKTPLIFVPQGIKVNGNNYLDMLKTELMPWVKKHFKKTKWTFQQDGAPAHKHKNVQAWCESNFPDFIAFNQWPPSS
PDLNPMDYSVWSVLEAKACSKPHRNIDSLKDSLKKAWDELDINYLRATVDSFPRRLEACVAANGDIFEL

**>DD37D CbmaT4 (AC084524 REGION: 10169..11203)**
MKSSPHRPSIELLFKRGLGSAEIARRLQISSSTVRNVVAAIKKRGDASEVKKSGRPRSVNTRNTRAIIKKRIIRNDGLSLNRMASQLGI
ARSTVQSIVKNDLKLKSYKLRRGQYLSDKSKAMRLEKCRKLLQHFQVRRVSDVIWTDEKIFTIEPLPNRQNQRQLLSKDDSMSPKRRLA
HNRLFPKSVMVWAGITATGKTPLVFIERNVKINSEVYQKIVLMDNLLPWVTQHFAGGPFILQQDWAPSHGSRSTLAVLEAHFPGFLDKN
LWPASSPDLNPMDFSVWGMLEGKIAGKVFATVDDLKAALEVAWASIDDGYLRRTVNSVKKRLRACVKARGSNFEFLL

**>DD37E Ae-atropalpus1 (AF377999)**
MEAERRDKIVHSFLENPLLPASKLAKQLKMPRNTVWRVIKQYKETLMTAPKPHSKRRSGTVDRKLRGKVIKAIKRNPNLSDRDLANKFQ
AAHSTVRRIRLREGRRSFRASKQLNRTLKQNNVARIRARKLYDQVLTKFDGCILMDDETYVKAEFGQIPGQKIYLATARGDVSAKFKFV
FADKFARKNIIWQGICSCGQKTKVFLTDKTMTSEVYKKECLQKRILPFIRAHDRPVMLWPDLASCHYSKTVIEWYATNGVSVIPKDLNP
PNCPQFRPIEKYWAITKRRLKAKGKLVKNMTQMKNWWNQIAKTVDEKGVRRLMCRITGKVREFLRNSNE

**>DD37E An-gambiae1 (AF378002)**
MEAERREKIVHNYLENPLWSASRLAKKLKFPRNTVWRVIKRYKEILTTIRKPQANRRSGTVDQNLRSKILKTIKGNPNLSDRDLARKFG
ATHSTVRRTRLREGIKSYRASKQSNRTIKQNSLIKTRARKLYDQVLTKFDGCLLMDDETYVKADFGQIPGQTFYLATGRGDVPAKFKFV
FADKFARKFMIWQGICSCGKKTKVFVTNKTMTSELYQKECLQKRILPFIRSHDHPVMFWPDLASCHYSKVVREWYAEKGVLFVPKNLNP
PNCPQFRPIEKYWAIMKRRLKAKGKVVKDINQMTTWWNKIAKTMDEEDVRRLMSRVKGKNREFLRNREE

**>DD37E PrDD37E1 (DQ138288 REGION: 3859..5197)**
MKDLPRSGRPFKLSKKKIDCLVESVNNRCGVSQRKLARRFGVHQSTICRNLRRRTSVVIRKRKKAPKMNSTEQETRAQENCGKLCRMLV
NGCDIIMDDEKYFKLSGDNVLGNRYFYSNDPSTAPPDVKFQKKAKFERKVMVWIAISSRGISSVYVHKSKQAVRQETYLAECIDKRLLP
FIEKHHSDGKYLFWPDLATSHYSNIVQQRLRDNHIPYVLRIDNPPNVPQARPIEQVWSLLEQKIYENNWEAKDIDCLARRIKQKVKEFD
QDMLRRMICSVRKKLLSMWKKGLYSVI

**>DD37E DrTRT (Zhang et al. 2016a)**
MGQKKDLTGSEKSKIVRYLAEGCSTLKIAKLLKRDHRTIKRFIQNSQQGRKKRVEKPRRKITAHELRKVKRAAAKMPLATSLAIFQSCN
ITGVPKSTRCAILRDMAKVRKAERRPPLNKTHKLKRQDWAKKYLKTDFSKVLWTDEMRVSLDGPDGWARGWIGKGQRAPVRLRRQQGGG
GVLVWAGIIKDELVGPFRVEDGVKLNSQSYCQFLEDTFFKQWYRKKSASFKKNMIFMQDNAPSHASKYSTAWLARKGIKEEKLMTWPPC
SPDLNPIENLWSIIKCEIYKEGKQYTSLNSVWEAVVAAARNVDGEQIRTLTESMDGRLLSVLAKKGGYIGH

**>DD37E CmpTRT (Zhang et al. 2016a)**
MGQKRDLTDSEKSKIVKSLSEGCSTLEIAKILGRDHRTIKRFVANSQQGRKKRVEKKRRTLTAKDLRRIKREATRNPLSSSAVIFQNCN
LPGVPRSTRCSVLRDMAKVRKAETRPPLNKTHKLKWQDWAKKYLKTDFSKVLWTDEMRVTLDGPDGWACGWISNGHRAPLRLRCQQGGG
GVLVWAGIIKDELVGPFRVEDGLKINSQTYCQFLEDTFFKQWYRKKSASFKKTMIFMQDNAPSHASKYSTAWLASKGLKDERIMTWPPS
SPDLNPIENLWALLKWEIYGEGKQYTSLNSVWEAVVAAAQNVDCQQIKKLTDSMNGRLMTVIEKKGGYIGH

**>DD37E TrTRT (Zhang et al. 2016a)**
MGQKRDLTGSEKSKIVRYLAEGCSTLKIAKLLKRDHRTIKRFIQNSQQGRKKRVEKPRRKITAHELRKVKRAAAKMPLATSFAIFQSCN
ITGVPKSTRCAILRDMAKVRKAEKRPPLNKTHKLKRQDWAKKYLKTDFSKVLWTDEMRVSLDGPDGWARGWIGKGQRAPVRLRRQQGGG
GVLVWAGIIKDELVGPFRVEDGVKLNSQSYCQFLEDTFFKQWYRKKSASFKKNMIFMQDNAPSHASKYSTAWLARKGLKEEKLMTWPPC
SPDLNPIENLWSLIKCEIYKEGKQYTSLNSVWEAVVAAARNVDREQIKTLTESMDGRLLSVLAKKGGYIGH

**>DD37E SsTRT (Zhang et al. 2016a)**
MGQKRDLTDSEKSKILKSISEECSTLEIAKILGRDHRTIKRFVANSQQGRKNRVEKKRRKLTAKDLRRIKREATRNPLSSSAVIFQNCN
LPGVPRSTRCSVLRDMAKVRKAETRPPLNKTHKLKRQDWAKKYLKTDFSKVLWTDEMRVTLDGPDGWARGWISNGHGAPLRLRRQQGGG
GVLVWGGIIKDELVGPFRVEDGLKLNSQTYCQFLEDTFFKQWYRKKSASFKKTMIFMQDNAPSHASKYSTAWLASKGLKDERIMTWPPS
SPDLNPIENLWSLLKRTIYGEGKQYTSLTSVWEAVVAAAQKVDGQQIKKLTDSMDGRLMTVIEKKGGYIGHRFFF

**>DD37E HbTRT (Zhang et al. 2016a)**
MGQKKDLTGSEKSKIVRYLAEGCSSLKIAKLLKRDHRTIKRFIQNSQQGRKKRVEKPRRKITAHELRKVKRAAAKMPLATSLAIFQSCN
ITGVPKSTRCAILRDMAKVRKAERRPPLNKTHKLKRQDWAKKYLKTGFSKVLWTDEMRVSLDGPDGWARGWIGKGQRAPVRLRRQQGGG
GVLVWAGIIKDELVGPFRVEDGVKLNSQSYCQFLEDTFFKQWYRKKSASFKKNMIFMQDNAPSHASKYSTAWLARKGIKEEKLMTWPPC
SPDLNPIENLWSIIKCEIYKEGKQYTSLNSVWEAVVAAARNVDGEQIKTLTESMDGRLLSVLAKKGGYIGR

**>DD39D Soymar1 (AF078934)**
MQRKVKMLSNEERITIYQLLLQKSVDGKLPQGVKESVASSFSVCRKTIDRIWKRAKESETHDVSHKKTKNSGRKRVEIDLSQLREIPLS
QRTTVRTLAVAMKTNTSAMYRLIQSGAIKRHSSAIKPQLTEEGKRLRLEFCLSMLEGIPHDPMFQSMYNIIHIDEKWFYMTKKSERYYL
LPDEDKPHRSCKSKNFVPKVMFLTAVARPRFDSEKNVTFSGKIGIFLFVTQEPAKRTSVNRVAGTMETKAITSINRDLIRSVFIEKVLP
ATKEVWPRDELGSTIFIQQDNARTHINPDDPEFVQAATQDGFDIRLMCQPPNSPDFNVLDLGFFSAIQSLHYKEAPKTIDELVNAVVKS
FENYCVVKSNFIFLSLQLCMIETMKAKGSNRYTSQHMQKEKLETEEQLPIQLKCDPILVQETLDYLNNN

**>DD39D Br-oleracea (XP013589454)**
MTDEERVEVYHALLERSNNGKLLKNSTREVSGLLDVPLQTVQNIWKRAKNTGYGEVVDVSHRRKGKCGRKKKQIDWLKVVDIPLHRRTT
IRSLAAALGMSPTVVFRSLKEGQLRRHSNAIKPLIKEDNKKTRVKFCLSMLNKNALPHQPKFVDMYNVVHIDEKWFYMTKKTQTYYLLP
SEEDPLRTCQSKNYISKVMFLAAMARPRYDGEGNETFSGKIGIFPFVTLQKAQRRSCNREAGTMELKPMVSIKREDIKHFLIEKVLPRI
YERWPAEDFGKTIFIQQDNAKTHVTVNDEEFQVAALQHGLDIQLMCQPPNSPDLNIFDLGYFRAIQALQHHVCPKTVEDLVTAVEEAYD
EYPPNLVNRVFLTLQSCMIEIMKIGGGNNYKIPHLKKDTLEREGLLPVQMDCDPNLVEEAMNYVAC

**>DD39D Ca-sativa (XP010462775)**
MTDGERLKVYHALLERSNNGTLKRTHTREVANLLSVPLLTVQRIWKLAKDIPNGEVVDVSHKRKGKCGGKNIVFDLDRIVDIPFNRRKT
LRSLAAALKISRTTLWRCLKRGLIKRHSNAIKPRLTENNMRARLQFCLSMLDRTTLLGHPKFVDMHNVVHIDEKWFYMTKRSENYYLHP
SEEEPYRTCQSKNYIGKVMFLAAMARPRFDNNGNETFSGKIGVFPFVTMQPAQRHSRNREAGTLELKPMTSIKRENINDFLIGKVLPRI
RERWPQEDFEKTIFIQQDNARTHVDPRDEDFRAASSHHGFDIRLMCQPPNSPDLNILDLGFFNAIQTLQHEVCPKTIEELVSAVEVMFD
EYPPYLVNRIFVTLQSCMQEIMKV

**>DD39D Phyllostachys edulis (ADP24264)**
MANLDLNQPIHWEEIEDYDGPVIDLNFDLVFHDSDEGEDGGPTHGEEDGTLAHGEEDGGAPSHGEEDGAPAHGEEDGAPAPNAYETIST
NKAKNCGRKWVAFDPEAIKDVPLSSRTTIRDLAGALNISKSTLFRRMKEGKFRRHTNDIKFTLTEDNKRARVKFCLSMLDKLSMPQEPT
FEGMYNIVYIDEKWFYRTRKCQNYYLALDEDKPERTTKSKNFIEKVMLLAAIARPRFDGDGNVTFSGKIGIFPFTFVEPAKRSSANRPA
GTLVTKAMTSVTKETSREYLVNKVLPAIKQKWPAEEVGTPIFIQQDNARTHIAINDDEFCRAASADGFDISLMCQPPNSPDLNVLDLGF
FAAIQSMFQKSSPSNVEDIVAKVIQAFDEYPVDRSNRIFLTHQSCMREILRQKGGQHYAIPHLKKQSLERNGVLPVSLQCDPEVVNEAI
VYIN

**>DD39D Pisum sativum (AAX51974)**
MYIYHELLQKSVDGKLRKGATNEVASSNSVPLRTVQRIWKRAKESETRDVSHRKTKNCGRKRISIDENQIRELPFSQRTNIRSLAFALK
TNPTSVFRLIKSGAIRRNSNAIKPLLKEENKISRLEFCLSMLEGTPHDPMFKSMHNIIHIDEKWFYMTKKSEKYYLLPDEDEPYRTCKS
KNFIAKVMFLVAQTRPRFDSEENETFSGKIGVFPFVTHEPAIRSSINRVAGTMVTKAITTVNRDVVRSFLIDKVLPAIREKWPRDEFES
TIFIQQDNARTHINHDDPLFREAATKDGFDIRLMCQPANSPDLNILDLGFFSAIQSLQYKEAPKTIDELISAVVKSFENFPSIKSNRIF
VSLQLCMIEIMKEKGSNKYKIPHVNKERLERVGQLPIQIKCDPILVQEVKNYLNME

**>DD41D Crmar2.5 (AAK61417)**
MERYTIQQRVKVIQTYYENGRSNQNAYRALRDFFGQFDRPNVRTIAKIVEKFEQIGSVEDVRTPVHARTARTAENIAAVRDSVAEEPST
STRRRAQQLHLSRSSMMNIMHKDLHLHAYKVQLAQELKPLDHSKRREWAEWFQEMATVDDQFSKKIIFSDEAHLHLSGFVNKQNCRIWA
NENPRVIVEKPVHPQRVTVWCGLWAGGIIGPYFFQNEAGQAVTVNGVRYREMITNFLWPQLEDMDVDDMWFQQDGATCHTANETMALLR
NKFNGRVISRNGDVNWPPRSCDLTPLDFFLWGYLKEKVYVDKPATTQELKDEIIRHINGIETPLCLSVIENLDHRMEVCRRGRGAHLAD
ILLHT

**>DD41D Apismar4.1 (Bouallegue et al. 2017)**
MERYSKEQRVLIVKTHYQNGEHYAVTVRKLRTILGHHNAPNESTVRRLIKKFEESGSTQDKKISGRHRSGRSEANVTVVHDSVTVSPRK
SCRRRAQEMHMSPATMQRILTKDLHLHAYKVQLTQELKPADHEKRRQFVEWILTRDRESEGFAKRIIFTDEAHFHLNGFVNKQNCRIWG
SENPRTIQEKEMHPERVTVWCGIWSGGLIGPYFFEDEEGNAVTVNGVRYRAMLNHFLWPRLDQMNIENVWFQQDGATCHTSRETIALLR
EKFPDTLISLRGDQSYPPRSCDLTPCDFFLWGYTKSRVYQNKVRNVLELKQEIRCVLNELDGAMCDRVMVNFMERIIAYRASRGGHMPD
VVFHC

**>DD41D Mariner-12_CGI(Repbase, https://www.girinst.org)**
MEPSCEQLAEIVKLYYECKSVTLVIRTMRKRHPEMKMLNRMLIYRLVRKFEGKGTISDTRNNSGIGRPRSVRTKENIDAIDRLITETPQ
KSVRRVLHELDCNVSKSSVHRILKFDLKLIPYKISVMQHLKETDIRSRLEFAHWILSRPDSENLTDAIWFSDEAHFHLNNVVNKQNFRF
WGSAKPNFYVEKPLNCEKVTVWAALSSTGIIGPFFFEDTEGEAVTVNSERYLKLMKSKFLPALRRKVVNFDDVWFQQDGATPHTAGCVL
QWLDNTFGDQYISYRTANVWPPHSPDLNPLDFFLWGYLKDRVYSPSPQTTEDLKTAITREIRSISVDTCKAVIKNFRNRVQTMLKAKGR
HLEHML

**>DD41D Lsra_Ap (Zhang et al. 2016b)**
MLVVILVRLVGLNVKMNNQEKVQMLLIYGKCDRNSRQSARMYAEQYPGRYHPPHTFFIKIEQLLINHGAFSVKVVRNQQIRENNINEDV
ELQVLAYIRLNPRSSVRHVGREVGISFGLVHKILKKHKMHPYKPDLVQHLRPADPERRLNFIAWLLVQIDTKPLFLNQILWTDESKFTN
NGVINKQNNRMWSDVNPHWAVDNRYQTVWGTNVWCGLIGGKLLGPYFYEENLTARRYLAFLTNVLPLMLENLPLATRQTLYFQQDGAPA
HNAHIVRDYLNRVYEGKWLGTYGPIEWPARSPDITPLDFFLWGHLKTVVYADPPVNLADLKNKILVACNNLTESQIMSATNRGCLQRFQ
LCVDNHGANFEQFI

**>DD41D rosa_Ae (Zhang et al. 2016b)**
MATPNELQTRNFDSLLVIPSAARNNMFSLEERFEILKTYFQSQCCVAETVRILKRNMGRDRAPTEGAIRKLVRKVREKGMLVDDRSGPR
ARTVRTPENIEAVAQSVRQNPTTSTRRRSQQLSISRTSLRRILHKDLGLFAYKIQMTQELKANDHPLRYQFAVWAIDQLNNDVDFGRKI
IFSDEAHFHLGGYVNKQNCRIWGSEQPHTIVERPMHPPRVTVWCGFWSGGIIGPFFFENDAGNAVTVNGERYRAMLTNFLWPQIDTMNV
DDLWFQQDGATCHTSRLTIDLLRTKFNNRIISRSGDVNWPARSCDLTPLDFFLWGAVKDRCYADNPETIPALKNNITTVLSEIGAEVVQ
NVINNWGDRMAYCKASRGSHMNEIVFHS

**>DDxD pogoR11 (S20478)**
MGKTKRVVGLTLKEKLQIIELVTNKVDKKEICAKFKCDRSTVNRILQKTNEIHEAVAASGLKRKRQRKGAHDLVEEALYIWFGQQESKN
VILDRHVILAKAKEFCQKFNDAFEPDASWLWRWRKRHNIKYGKIHGETATNDSVSANEYKNDILPGLLKGYNPEDIFNADETALFYKAM
PNATFFTCGKQLNGQKSQRVRLTLLFICNATGTYKKTFVIGRSKSPRCFKNANVPIPYYANKKAWMTKDLWRKIMTGFDEEMKKQNRKI
LLFIDNATSHTTVKDFENIKLCFMPPNATALLQPLDQGIIHSFKLEYRRILVKQQLIAVNCGKSTVEFLKSLSLLDALYFVNQGWKNVK
MLTIQNCFKKAGFKFSFENEDTIAEKDKQCVEVDIVSNINWNEYANVDADEACHGQLDDDEIVRSLVQDAKTSDNEESHSDEDVDDTER
PTFKDGFAAIKALKSIFMRNNNDEFLQNLNSMEDKLFNLHINSAVLQKKITDYF

**>DDxD Tigger1 (U49973)**
MASKCSSERKSRTSLTLNQKLEMIKLSEEGMSKAEIGRKLGLLRQTVSQVVNAKEKFLKEIKSATPVNTRMIRKRNSLIADMEKVLVVW
IEDQTSHNIPLSQSLIQSKALTLFNSMKAERGEEAAEEKLEASRGWFMRFKERSRLHNIKVQGEAASADGEAAASYPEDLAKIIDEGGY
TKQQIFNVDETAFYWKKMPSRTFIAREEKSMPGFKASKDRLTLLLGANAAGDFKLKPMLIYHSENPRALKNYAKSTLPVLYKWNNKAWM
TAHLFTAWFTEYFKPTVETYCSEKKISFKILLLIDNAPGHPRALMEMYKEINVVFMPANTTSILQPMDQGVISTFKSYYLRNTFRKAIA
AIDSDSSDGSGQSKLKTFWKGFTILDAIKNIRDSWEEVKISTLTGVWKKLIPTLMDDFEGFKTSVEEVTADVVEIARELELEVEPEDVT
ELLQSHDKT

**>DDxD Fot1 (Q00832)**
MPVYSADDLENAIADFKNGVSLKTAAKKNGLPPSTLRGRLTGAQSRQVARQEQLRLTTDQEDDLERWILRQEKLGHAPTHAQVRTIVRS
VLARHGDHAPLGRKWTTRFVERHPALKTKLGRRTDWERVNAATPANIKRLFDVYETVDWIPPERRYNADEGGIMEGQGVNGLVIGSSQE
SPNAVPVKTATVRTWTSIIECISAVGVVLHPLVIFKAKTIQEQWFRREFLQKHLGWQVTFSKNGWTSNSIALEWLEKVFLPQTAPADPA
DARLLIVDGHGSHATEQFMAKCYLNNVYLLFLPAHCSHVLQPLDLGCFSSLKAAYRTLVGEHTALTDSTRVGKQRFLDFYARAREIGFR
KVNIRSGWRAAGLWPVNINKPLASRWVMVLTKSALPPSETLDIATPKRGGDVVKLFSAKSSSPSSRLSIRKAAAALDKVAIELAMKDRE
IERLRAQLEAAQPKKKRKIRQDPNECFISLAQILAEANREPDQRVIQSQKGDLDCIVVDGKSSSESEEDPAPVRRSTRVRRATKMYIRQ
DLSSEESD

**>DDxD Tan1 (U58946)**
MPPKASIPSKSQVEREGRILLAIEAIRKGQITSIREAARVYDVARTTLQARLSGRVFAKNMTNARQKLSNNEEESLVKWILSLDKRGAS
PRPLDIRDMANLIISKRGYSTVEQVGINWAYSFVKRHESLRTRFARRLNYQRAKMEDPEVIKDWFKRVQEVIQEYGISSDDIYNFDETG
FAMGMIATYKVVTSSQRAGRPSLVQPGNREWVTAIECIRSNGEVLPSTLIFKGKTHLKAWYEGQSIPPTWRFEVSDNGWTTDKIGLRWL
QKHFIPLIRGKSVGKYSLLVLDGHGSHLTPEFDQSCAENEVIPICMPAHSSHLLQPLDVGCFSVLKRTYGGMVQKQMQYGRNHIDKLDF
LEVYPKAHQCALSKSNIISGFRATGLVPLDPDQVLSRLHIRLKTPPTPDSQSSGSVLQTPHNIKHLLKHPKSVERLLRKRQASPTSPTN
STLRQLLKGCELAITNSIILAKENAELRASHEKQLPKRKRSRKQVIYTEGTTVEEAQRAIQEVEEVQNDEDIEVEPQSQYTETPSRAPP
RCSNCFNIGHRRTQCSKPPTN

**>DDxD Pot2 (Z33638)**
MKQYTEKQLISAINDVNNGNPIAKTSRKWGIPRSTLQSRLKGSQPYKKAQSPFQRLSTEQEKHLADWVLTQTALGLPPTHQELRFFAER
ILQAAGETKGLGKRWITRFLARYPILKTQRPRRIDNARVNGATTEVIKSWWLYITNPVINAIKPENRWNMDETGIMEGKGSNGLVLGLN
GIRPLQRKEPGTRGWTTIIECISATGVALPPLVIFKGKNVQQQWFPTDLSPFDNWQFHATENGWTNNQTAIEWLKKVFIPYTQPLTPEK
RLLVLDGHGSHITDEFMLLCLQNNIQLLYLPPHSSHVLQPLDLSVFGPLKEAYRRQLGFVSQFCCSTVIGKRNFLLCYRKARLKAFIAK
TIQSGWRTTGLWPVNLVKPLLSPFLLENSNANVIKDKNNGLQRDKTPESPAQKINDPSLLIWKTPKTTRDIRLQLQKLSQSNKTNATSR
LLFAKVQKSFEAKDTLLASAQQKISLLEAQLEAIRPVKRRRVVPDPNELLVNKQNIIGLQENDIENLEPLADEEEVNEPEKRENDCIFV
R

**>DDxE RS(alfa) (X02581)**
MIPEAREVHLSRKDRKVLEAVCRSPVTLQRDLKRARIVLLAADGRSTRSIAKEVGVQPRIVSLWRHRYADHGLEGLQDKPRPGKQPIYT
KTTDKRILKLLDKPPPQGFARWTGPLLAEALGDVDVQYVWRFLRSHKIDLVARKSWCESNDPNFTAKAADVVGLYVAPPAKAIVLCVDE
KPSIQALERAQGYLKLPSGRALTGQSHDYKRHGTTTLFAALEVATGKIIATHSKRRRRVEFLDFMNSVTAAFPNRKLHVILDNLNTHKK
NEDWLKAHPNVQFHFTPTSAPWLNQVEVWFSILQGQSLSGTSFTSLKQLQEHIDAYVNAYNDRAEPFVWTKKKVRQRRFKGRRRITQL

**>DDxE IS630Ss (X05955)**
MPIIAPISRDERRLMQKAIHKTHDKNYARRLTAMLMLHRGDRVSDVARTLCCARSSVGRWINWFTQSGVEGLKSLPAGRARRWPFEHIC
TLLRELVKHSPGDFGYQRSRWSTELLAIKINEITGCQLNAGTVRRWLPSAGIVWRRAAPTLRIRDPHKDEKMAAIHKALDECSAEHPVF
YEDEVDIHLNPKIGADWQLRGQQKRVVTPGQNEKYYLAGALHSGTGKVSCVGGNSKSSALFISLLKRLKATYRRAKTITLIVDNYIIHK
SRETQSWLKENPKFRVIYQPVYSPWVNHVERLWQALHDTITRNHQCSSMWQLLKKVRHFMETVSPFPGGKHGLAKV

**>DDxE IS630Se (NP_073225)**
MPIIAAIPDEERQLMRKEAQQTHDKNHARRLIAMLMLHQGMTVTDVARLLCAARSSVGRWINWFTLHGVEGLKSLRPGRAPRWPVADIL
QLLPLLVQRSPKDFGWLRSRWSTELLALVINRLFDVTLHRSTLHRYLRQADMVWRRAAPTLKIKDPHYEEKRLVIDQALAQEQTAHPVF
YQDEVDIDLNPKIGADWMPKGQQKRIATPGQNQKHYLAGALHSGTGRVHYVSGSSKSSDLFISLLETLRRTYRRAKTITLVADNYIIHK
SRKVERWLEENPKFRLLFLPMYSPWLNPIERLWLSLHETITRNHQCRYMWQLLKQVAQFMNAASLFPGNQQGLAKVER

**>Mariner-18_CGi (Repbase, https://www.girinst.org)**
MSITSSNKMTSSVKKQTSCTSVDTVNERAQALALLDAGISVKDVAKQLCKSERWVTKWRRRREQNKQLTDQPRSGRPSKIVGVVKRKVE
NIKYKRGKSTRKCSKELKNSGFNVSHVTVFNYLRKVKKWKAFKRSRNQLLTKTQRQKRLKFARDHKHLTAEDWENYIFSDESSKYLFHV
PNRQDDVVWGSQSDEVPDVSCVKSSAKVMIWGAMGVNGLSKLHIVPSGKTINAKYYVDKILQKELKPALNRQKNTGRIDERKLVQYPGH
ATFVQDGATPHTALVTQNWCKDNLPNFISKEEWPGNSPDLNCIENLWSILDSEAYKDPRPTSMDQLRRRLQRAWREIPQKHLISLIHSM
PNRIKNVLKNKGGLSGY

**>DD37E(L18) Nematostella vectensis (NW_001834331 REGION: 451691..452376)**
LRCQQTNSKKNSKNSRKKQEFGFKVYSSLAKFSRQAKKWNAESAKSSCKMVVENAKRVQERIFIQKKKNILKRKAILGFLLTVWRYMTR
KGWKPFKQQKKPLLSTQQKKKPFEVCTEVQISVRF*MGNFLFTDECPKYLLYLPN*KNNVV*GSQETEVPPSWHDQPGSMTGRGLTSLH
FLPQGRTVNSEYYIKRILEKGVKPFSRKSNSENLAKRMLFSNERFMVFVQDGAPAHTTKASQHWCKKNLPASLEKAEWPANSPDLNPIE
NLWGIIDEETYSDPQPKTMTSLKSRLKECVIVSVPRMCHCLL

**>DD37E(L18) Apostichopus parvimensis (JXUT01105677 REGION: 368..1513)**
MAGKKGEQASANERKRVVALRTVLDSFSTVLDRYSFREVATMLNRSEGFVKKQWARRHWPSLNNKKRESRSRKLTPEAKRFITNFVKDK
WSRGYRKAAKALNSSGLLGNGVSVNPTTVLRYIKSQPWGKTAYKVEVKPLRTGNQVEKRLAACNYWNRQGFLDATEEGRKKRMNIMWTD
EKNFGLFHKPNKQNMRMRTEHKSSVPVARIVKDPLKIMVSGGMTGYGVTDLLIMEPKTRVNGEFYRQKMINDIFAPAFNKTEGPDKLFD
RTANPIFMQDGATSHTAKLTQELCSRVFPNYWDKNFWPGNSPDMNPIENLWSIMEAKVYCPPLATNKNMLVKKIQDAWVYLKLRKHEIL
PNLLEGDNGWRSRCQKCIENYGGDSS

**>DD37E(L18) Hydra magnipapillata (EQ256867 REGION: 98791..99915)**
MGKVIGKSNLAASLKKRIAELGKLGKSKAEVAAAIGCSEKSIQRYWRKPVNTNFNEKKRCGRPTVLSPASKNLITSEMKDKWGSSTRSC
AKKLNFSERYITRKKQISRSTVQRFVQHQPWGKVAYHKPIKPLLTEKNQNDRLKFRDWLEQNGYLQDEHVGRQKRGHVLWTDESPVELF
PVPNRQNMRIWTDDKSKITPAVCPKFGLKIMVCGGMSRYGLTELVVVPEKQTVDADYYINHILPKYVEATTRNHQGDTADKRKMFFNQD
MILFQQDGAPAHSAKIVQQFCAKNFPSMIPKELWPGNSPDINVIEHLWNFLQQSVFEAPKPKNRVELVERVKDKWSSVTGDYLCLLVKS
LPKRVQEIRAANGGHSSY

**>DD37E(L18) Amphiprion ocellaris (NXFZ01003443 REGION: 185337..186248)**
SLKKKAQALGKNIHWVQKWWQRCNKGGSFKDLPRSGRPSGLTARVKDLMRKTKSKRHQSCR*LSQKLKNLGDVSKSTVHHYLTEPLELN
AYKGQIPYDSQKRFAFTKKSITLTVKDWETWISSDECPLYLFLTPSSQNNRIYTDDHEKVPSFEQVKFSTHCKV*GSVSASGLSELHIV
PQGTTVNARYYTNNILEKVLLPVLARRKKSGPVTEQKMVNVCSELVFMQDGAPAHTARMTQQWCSEDLPGFLHKEEWPPNSPDLNPIEH
CWGILNSHIFTSPPSTKKQINSRSARESGKIESVLKD

**>DD37E(L31) Crassostrea virginica (MWPT03000010 REGION: 5284904..5285974)**
MAKINSAGRVYDKGTPLGIDLRRSIINYMEEQGAKFGSLQLPRGLPQRVSEIFKVSHPLVTKIWKQYCIEGIVKLPEYKSGRKRKLDQE
DVQHIHFLKETKPSMPLKSVKEEVLKYSNAVIKVSESTISRHLKNDLNMTYKRIARYSKNRFTPQNMNYTQNFLNYVGQKDPFSLKFMD
EMGVKLSDGQNVYGHSLKGTPCIEMTRHNPHRNVTASVIVGISGVKYVKIFDGASNGTEYTQFIAEATQSYTDEGEPVFNPGDCLIADN
APIHHNMAERELNNYLPTVGVEYVFLPTYSPDLNPAEQVFRKVKKILKCDRYITLLNLDLKVAVYEAFKEISTADTLSFFRSTEYIKNP

**>DD38E(L31) Mizuhopecten yessoensis (NEDP02003490 REGION: 263291..264307)**
MEVTQSGRYFTKGERITDAEKTKVIELYSAGLPYSQIAKSTGITKGCCFKITQTFTTDKPQRDPVISPKLTNEVLQFIE*QKIAKPSIY
AKEIYVKLLETNVCTMNNIPSVRTIHHALKSILGMTHKVLQRIPSETTTDQFENKLNRFLT*LLLYTPEQLHFFDEASIVRTDGNRKRG
HNYRGEKAVEIQKYASNATFTINLCTWYFGIDHFGIIEGSSNANIMLKFFDEAMQEMNIVGNPVLALGDCIVIDNCGFHHQRFGEAFLR
HMLGIRGINLVFLPPYSPELNPCEYVFKLMRYRLRQNTALTYEYTEYAVVNAVTGIPDCIMPKLFTNCGYV

**>DD37E(L31) Mytilus galloprovincialis (LNJA010281983 REGION: 974..1891)**
IKLLNKLCIIHVSGKKYNENLSVSRRP*AGGRPRKYGIDEIEFVNVLKTERPSVETEHFKRPAPSVFCH*QHLYKYSIKNNKKNDLRMT
YKRIAHYKKNRFTVRNLQYTQQFLNYVSNKDPFTLKFMDEMGVKLVDGQPVYGHSRKGTPCVEITRYDPHANFTASLIIGITGVKYVKI
IEGASDSGEYLQFIGEASQSYTNDGESVFQPGECLVVDNEPTHHNMSERVLRNWLPTVGMEYLFLPAYSPDLKPAEQCFRKVKKLLKSD
RFGPVLRQDLKVAVYKAFNKITLIDTRSFFKANEYMNI

**>DD37E(L31) Exaiptasia pallida (LJWW01000515 REGION: 97496..98524)**
MAAVNSLQRPYTPGKPLSIAERQHIIELFNGGLTKTKVSERLRVTFRCVTNVIEHYRRYGSANPLGHSGKQPVVLTDDILEVVEVWKHQ
KPSLYASEIKDRLILEGICHWSTAPXVSAINRAXSTKLDMXWKKITSVPSEYYNNEYKVDDYLEITSRLDPSTLXFFDESSVIKTTSNR
LYGSSFKGYRAIEIQRYASNATFTVNLLHSILGVDYYNVIPGASNGEELVAFFSYALDCERQNGLPVFMNGDTVIMDNCGFHHGHITEQ
ALRHILGAKGVNLVYQPPYSPHLNTCEYCFNQMKQTLKLDEHFSQAYTELAIMHAVNNITPTQSLNYFKKCGYVF

**>DD37E(L31) Acropora digitifera (BACK02007470 REGION: 11157..12218)**
MSKVNKCGRTYNPGEAIGESLRASIIDRMLLDGGDPATGFFGGRFKNIGDCFGVSAPFVSKLWKTFCLQGDHMPQKRSSGNPSHLKPED
VQLVEFLKKEKPSATLASIKETVENYCNLNGGTSLSAIGNTVRNRLPDGPFTRKKLTRPKAEKFTPQNLAYCGQFLNFISALPPEKIKF
FDEAGVNSGTGNPVYGSSLKGTKSIEIAAIPRGVNITLNLLVGLEGILYANTLDGASNSFTFLNFFGEAGQATSALGNPAIEQGDFIIM
DNCAIHRFEAGTALQTWLMDMGANVIYTPSLSPEFNAVELVFNKLKILLKREEYVPLLKENVHVAIYESLDQIDTNDMYGFYSLIL
