## Supplementary material for "Why did the *Tc1*-like elements of mollusks acquired the spliceosomal introns?": Online Resource 2

**Online Resource 2.** The features of TLEWI elements. Only copies with a query cover of over 10% were taken into account when counting the number of copies. Homologies for introns only are ignored. The copies with query cover over 95% were counted as the full-length ones. Reference transcribed RNA sequence is indicated by red letters. For the *Mytilus galloprovincialis*, two WGS projects were presented: APJB and LNJA. The results for both WGS projects are shown through slash (APJB/LNJA). TIR – terminal inverted repeats. Tnp – transposase. ORF – open reading frame. bp – base pair. aa – amino acid. "ND" – no data. "-" – not detected

| Species | Elements | Length, bp | TIR, bp | Tnp, aa | Signature | ORF | Copies (full) | Transcribed RNA sequence | Reference representatives localization |
|---|---|---|---|---|---|---|---|---|---|
| *Bathymodiolus platifrons* | *TLEWI-1_BPl* reconstructed | 3270 | - | 350 | D87D36E | 152..751, 1216..1365, 2481..2780 | 6(0) | - | MJUT01016913 (65168..68040) MJUT01045212 (464031..464800) |
| *Crassostrea angulata* | *TLEWI-1_CAn* | ND | ND | >309 | D87D36E | ND | ND | **GGIZ01057891** GGIZ01064667 GGIZ01034089 GGIZ01060883 JU024018 | ND |
| *Crassostrea gigas* | *TLEWI-1_CGi (Mariner-35_CGi)* | 3749 | 30/- | 351 | D87D36E | 415..837, 2076..2255, 2508..2657, 2784..3083 | 9(2) | **GECI01017794** GECI01029272 | NW_011935320 (101378..105126) |
| | *TLEWI-2_CGi* | 2160 | - | 354 | D87D36E | 308..739, 942..1118, 1331..1480, 1585..1884 | 1(1) | **GECI01015139** GECI01031748 | NW_011937980 (140700..142859) |
| | *TLEWI-3_CGi* | 5498 | - | 378 | D89D36E | 343..735, 1072..1251, 1527..1682, 2170..2436, | 3(1) | **GECI01033235** GECI01012373 GECI01007772 | NW_011937840 (1066254..1069133) |

| | | | | | | | | | |
|---|---|---|---|---|---|---|---|---|---|
| | | | | | | 4741..4878 | | | |
| | *TLEWI-4_CGi*<br>reconstructed | 2849 | - | 350 | D87D36E | 370..783,<br>1186..1371,<br>1471..1620,<br>1817..2116 | 2(0) | GECI01030859<br>GECI01037938 | NW_011937084<br>(1564..3453)<br>NW_011937070<br>(10802..12510) |
| | *TLEWI-5_CGi*<br>*reconstructed* | 2724 | - | 351 | D87D36E | 411..602,<br>1268..1828,<br>2114..2413 | 1(1) | **GECI01009412**<br>GECI01011883 | NW_011935478<br>(324240..327831) |
| *Crassostrea virginica* | *TLEWI-1_CVi* | 2798 | - | 345 | D82D36E | 280..699,<br>1567..1659,<br>1661..1732,<br>1845..1994,<br>2114..2413 | 249(1)<br>-3 | No TSA data | NC_035787<br>(41647301..41650096) |
| | *TLEWI-2_CVi* | 3613 | - | 349 | D86D36E | 348..767,<br>1553..1732,<br>2116..2223,<br>2226..2264,<br>2708..3007 | 12(1) | No TSA data | NC_035787<br>(13927588..13931200) |
| | *TLEWI-3_CVi* | 2984 | - | >342 | D87D36E | 139..321,<br>323..535,<br>1114..1293,<br>1585..1734,<br>2505..2804 | 2(2) | No TSA data | NC_035786<br>(9597070..9600053) |
| *Lithophaga lithophaga* | *TLEWI-1_LLi* | ND | ND | ND | ND | ND | ND | GFXL01009423<br>GFXL01018663<br>GFXL01034780 | ND |
| *Mizuhopecten yessoensis* | *TLEWI-1_MYe* | 5810 | - | 350 | D88D36E | 328..927,<br>1403..1552,<br>5339..5638 | 15(1)<br>12 | No TSA data | NW_018405537<br>(292996..298805) |
| | *TLEWI-2_MYe* | 1885 | - | >131 | D?D36E | ..,<br>323..415, | 3(0) | No TSA data | NW_018480030<br>(410132..412016) |

| | | | | | | | | | |
|---|---|---|---|---|---|---|---|---|---|
| | | | | | | 1210..1509 | | | |
| | *TLEWI-3_MYe* | 4409 | - | >279 | D87D36E | .., 1726..2112, 2630..2779, 3139..3438 | 3(0) | No TSA data | NW_018408611 (35526..39934) |
| | *TLEWI-4_MYe* | 4640 | - | 322 | D81D36E | 327..482, 1162..1359, 2213..2392, 2718..2783, 2785..2850, 4126..4425 | 3(1) | No TSA data | NW_018406970 (578599..583238) |
| | *TLEWI-5_MYe* | 4320 | - | >251 | D65D36E | .., 889..1119, 1122..1240, 3382..3471, 3474..3529, 3718..4017 | 2(0) | No TSA data | NW_018407999 (760561..764880) |
| *Modiolus philippinarum* | *TLEWI-1_MPh* | 6757 | - | 354 | D87D36E | 276..815, 817..888, 3287..3436, 5797..6096 | 9(2) | No TSA data | MJUU01046008 (66061..72817) |
| | *TLEWI-2_MPh* | 11460 | - | 350 | D87D36E | 294..893, 7739..7888, 9387..9686 | 5(1) | No TSA data | MJUU01070107 (57493..68952) |
| | *TLEWI-3_MPh* reconstructed | 9475 | - | 349 | D87D36E | 1311..1907, 6504..6653, 8518..8817 | 3(1) | No TSA data | MJUU01056176 (92051..106540) |
| *Mytilus californianus* | *TLEWI-1_MCa* | ND | ND | ND | ND | ND | ND | GFII01026741 GFIB01058122 GFII01024380 | ND |
| *Mytilus* | *TLEWI-1_MGall* | 5889 | 35/35 | 349 | D87D36E | 199..789, | 9(1)/5(0) | GFZN01000761 | APJB010606582 |

| | | | | | | | | | |
|---|---|---|---|---|---|---|---|---|---|
| *galloprovincialis* | | | | | | 1924..2073, 3431..3730 | | GFZN01078562 | (1304..7192) |
| *Nodipecten subnodosus* | *TLEWI-1_NSu* | ND | ND | >343 | D88D36E | ND | ND | **GFNL01233336** | ND |
| *Panopea globosa* | *TLEWI-1_PGl* | ND | ND | ND | ND | ND | ND | GFXE01013766 GFXE01212001 | ND |
| *Pecten maximus* | *TLEWI-1_PMax* | ND | ND | 349 | D87D36E | ND | ND | **GAOX01009905** | ND |
| *Pinctada martensii* | *TLEWI-1_PMa* | 4944 | - | 351 | D87D36E | 300..725, 2148..2324, 3582..3731, 4319..4618 | 3(2) | No TSA data | NIJJ01056843 (19675..24618) |
| | *TLEWI-2_PMa* | 2367 | - | >151 | D?D36E | .., 832..879, 881..985, 1734..2033 | 3(0) | No TSA data | NIJJ01083462 (26182..28548) |
| | *TLEWI-3_PMa* | 7563 | - | 348 | D88D36E | 376..789, 2431..2610, 5441..5590, 6807..7106 | 4(1) | No TSA data | NIJJ01019696 (5876..13438) |
| | *TLEWI-4_PMa* | 6826 | - | 345 | D87D36E | 586..741, 1958..2002, 2494..2700, 4112..4288, 5375..5524, 6090..6389 | 1(1) | No TSA data | NIJJ01022083 (23981..30806) |
| | *TLEWI-5_PMa* | 2400 | - | >209 | D87D36E | .., 468..644, 1299..1450, 1756..2055 | 1(0) | No TSA data | NIJJ01031006 (1..2380) |
