## Supplementary material for "Why did the *Tc1*-like elements of mollusks acquired the spliceosomal introns?": Online Resource 3

```
                                   *                *                                                   *     *
TLEWI-1_BPl     MAMGKVNR---LSISTRHLIIEYSKKGLSAVHIQRLLHRKYNITTTRQSIFMFVKRYSSTGVLAP-ATRRDNKFPRKLTDFQRRCIDMWLRHNSELTSQA
TLEWI-1_CGi     MAEQRGAR---LSVETKRRILAIRKEERSLGKIRSVLCLRHGVKVSRQGIYMFLKKWDRDQTLVR-KTRVPGIGQTKLKEIHKDLIQMWQMENDELTCGE
TLEWI-2_CGi     MAKMCKGG--RLTVRLKLKILDLYERNLSVLAIKDRLQKEDSVKFSRTTIYSFLKRHKSCGILTR-RRRTVNPASVKLKELHLKFIDLWMDKNNELKASE
TLEWI-3_CGi     MAPMR--------LDVKAQILKYREEGRGVLEICKHLRSHHGCTVSRQAIHRFLRH----GSLVR-RRKLVTNSARKILAIHRRFIHMWLTQNNELTACD
TLEWI-4_CGi     MVVNTRGR--RMSEGIKMMIVRYLLLDLYVEVIREKLKHLHGFVISRQGLYAFKRRWQTGKELRI--TRKICNGAVKLKKIHLKFMDMWLAKNNELTTGM
TLEWI-5_CGi     MAPGRGGR--SLSLSIRNIIIKLHKAGISPVKICKTLQDKHQFKTTRQSVRRFIIRFETTGCVHD-KRKPRRPEHTKVRRIHMQFINMWMAQNPEMTAAN
TLEWI-1_CVi     MALNSRGR--RMSEKIKMVIVQYLLMHLSVEAIRPKLKNFHSYVISRQGLYYFIKQWRSGKGLRR--TRNNSGNSVKLKSIHLKFMDMWLAKNNELTTEM
TLEWI-2_CVi     MAASKGTR---ILMETKKRIILLYNEKKSAGQIQKVLLQKYGERVSRQGVYMFLKRWKEDQVLTR--KKRGKSGNVKIKKVHKDLIQMWQRRNDEVTASD
TLEWI-1_MGall   MALRKSSN--RLSTVTRQIILKYAPK-FSALKISRLLEEKYDVKTSRQSVWRFLNRFKKSKSIRD-PPRRRVRG---ISDLHIKAIDQWLKKDNELTANA
TLEWI-1_MPh     MRSTRSAKVFRLSIVVRNLILKYAAAYFSTVKIQKLLQTNHQVITTRHAIRKFLDRYKITGCVSDIRPKNIAKSTSKVTVFLKQMIDMWYSQNPEQTSRM
TLEWI-2_MPh     MATSVSAR---LTIPSRKLIVDYKSRGYSTVKITRVLKEKHGIITTRQSVRLFLLRFKRTGSLRD-NRVNQYKQNAKRTAFLLMLLNLWLSQNNEMTSAA
TLEWI-3_MPh     MATARTSA-FRLSREARQIIIHYSSK-YSALKIGRILKEKYAITTTRQSVWRFLKRYQRTRNISD---IHRSAKPKRMTEFLVKTIDLWMYENTELTSNA
TLEWI-1_MYe     MAALQKIS--RLSPELRTSIIRYSDKGYAIRKIRKTLEETNNVTVTRKTVRLCIHRYRETGSIND---RQRTGAHLKITTGVRKVIQSWLSKNNELTSAD
TLEWI-4_MYe     MADLRRRR---LSVEAKLKIMSMKSK--TGGQITKILKEKYGILTTRKTVNIYRKRL--------------------VKPIHKKIINMWLSRNSELTSPE
TLEWI-1_PMa     MAATKRST--RLTTRLKAKILDLRKK-MSIQKVKETLEAEDNINISRQGLYAFLNANRTRSSLGR-KPRGVMPNHFKLKPVHLKVMDSWINNNPELTASK
TLEWI-3_PMa     MASSDRGK--RLSVETRKFILARKSDGLSVQRIKSELKKCLLIDVSRQTIYNIVSH----GSLVR-RRSTNGGNATKIRPVHMRFLNIWLSKNNELTSKD
TLEWI-4_PMa     --MASTGR--RLTITARKLILRYKEEGLSHAKIKQVLRDRHLITTTKKSIYRILAHFELVIC-----PLTNTYNSYKLTDLHRQVLHMWMQEKPESTSTA
TLEWI-1_PMax    MADPRSRR---ITMEAKLKIVSMKTK--SGSEIARILKENFNILTTKKTVNVYRRRIKRTGMLEIVRSSPALKNRQLVKPLHKKIINMWLSRNSEMTSPE
                                  PAI                                                          RED
                                     *                 **  *   *             *      **   **
TLEWI-1_BPl     LVDRLFRVFDVRVKTSYMSKVRKA-LGWCTRTL-QYCQLISHTNKLCRLQWSLDALRSKETFDNVIFTDETSVEMGADGGAFFYKRTS-DLDFLPAKKMK
TLEWI-1_CGi     MREKLKDSVGLDVSPALISKVRKD-LGWQAKTGNRTCQLISRKNVRERLQWCLKAVEDKEDFKNVIFTDETTVEMCSAGRLHFFQKSS-EIQKKTSRRSR
TLEWI-2_CGi     LQKRLFETFGLKVSTSLISTRRHSVLGWKSQTSNKTCQMISRKNKTVRMQWCLDVLQTRERFRDVIFVDESCVEMCANGRIAFHQKSS-DFEKKCNRIPR
TLEWI-3_CGi     IQRKLKDLFRLSVSLSCVRKVRRE-LGWSSNTG-KYCQQISHKNKKARMEWACNAIKNKDNFRNVVFADETSVEMCSHGKLFFHQSRS-GIEMKTRKRSR
TLEWI-4_CGi     LKRKLFEVFGVNVSTSLIRKRRSE-LGWNTVVS-KTCKLISKKNIQVRQSWCLDALNNKEDFRNLIFVDENTVEMSFSGRLFFHQPGS-KIQKICARRPT
TLEWI-5_CGi     IQGRLEKQCGIQLSKEYVCMLRRK-LNWTPKHM-KYGQLISHKNVKHRLDWCLDQLISKDSFRDVIYVDESTVEMCSSGRLFFHRHGS-NMDRLPAKAPK
TLEWI-1_CVi     LKRKLFEVFGINVSPSLIRKRRSE-LGWKTVVS-KTCQLISHKNKKVRKVWCTDALKNREDFRNVVFVDESTVEMSS-----FHQASS-LMQKTCARRPK
TLEWI-2_CVi     LQMKLLDSTGLNVSVQSIRKVRRD-LGFEAKSAGRTCQLISRKNVRERLTWCLNAVDKKEDFRNVIFTDETTVEMCSTGRLFFHQKSS-DIQQKTSRRSR
TLEWI-1_MGall   IADKLRETFSTNISKGYVSLIRRK-LGWTAKRV-KYCQLISHVNKEKRLHYAMKVLSMKEKFLNVIFVDESTVEMTSNGRIFFYKPDS-DSSQLPARKQK
TLEWI-1_MPh     IRNRLKDLFNVDVSNRLVSRIRRE-LGWTTRRA-KYCQLISNSNKLKRLNWCLKTLSSKDTFHNVIFVDETTVEMSSNGKLFFYKPNS-NLQRLPARKQK
TLEWI-2_MPh     ICHKFLTMFGMKVSTDYVCRLRKR-LGWTAQRI-KYCQLISHKNKHKRMEYALDCLNKSETFDNVIFVDETTVEMTNTGRLFFFKKGS-DLQKLPAKKQK
TLEWI-3_MPh     IAERLRHTFDLSFSKGHISRIRRR-IGWTAQRW-KYCQLISHVNREKRFQYAVKMMHSKEKFENAIFVDETTVEMCSSGRLFFYKPDS-NFSQLPAKKQK
TLEWI-1_MYe     ICARLKTELNLSVSRATINRVRVD-LGWVTKRL-QYCQLISHKNKRVRMKWSLDALARNDTFADVIFTDETTVEMNSSGRLFFHNTKCGEMQRVSTKKQK
TLEWI-4_MYe     IQRKFHSSFGLTVSTAHTRRIRRE-LGWTARHI-KYCQLISNKNKN-RYNFCINTLISLETFSNVIFVDETSVEMSSNGRMFFTQHGP-SMDRLPSKAPK
TLEWI-1_PMa     IKFKLKETFDLDISISLISLRRRH-LGWTARTSARACQLISHKNKKVRMQWCLDALQSRDDFKDVIFVDESSVEMTSNGRLFFYKPSS-DIQKKCDRKPK
TLEWI-3_PMa     IVDKLRSTFGVGISHGYARIIRRK-LGWMKSTG-KYCQLISNKNKAVRKAWCCDAWHRRDTFEDVIFCDESSIEMNSNGRLFFHRERSAGFSKTTTKRSK
TLEWI-4_PMa     LSLRFKEELGINVSQSHVRLIRRQ-MG*KTSQA-KYCQLISAPNRRVRLEWALNALSSNEKFEDVIFMDETTVEMSSTGRVFSHETVG-RLDCMANKQAK
TLEWI-1_PMax    LQRKFHSTFGFSVSTAHLRRIRKE-LGWKSRHV-KYCQLISNKNKINRYNFCIEALINLETFSNVIFVDETTVEMSSNGRMFFTQHGP-SMNRLPSKASK
                                                      NLS
                *** * *      *    *  * * * * *     *                   * ***** ** * **  *              ***** * *** *
TLEWI-1_BPl     PKHAYKVHV*SGISYRGRTSICIFSGIMDSVIFQNILQSNLLPFVEH--QFPDGFRLYQDNDSKHVSKSTKKWMEEHGILDKVMTTPASSPDINPIENLW
TLEWI-1_CGi     PKHAYKVHVWGGISYEGRTDICMFTGTMDSVIYQGIIEKNLIPFAEK--KFPNGFRIYQDNDSKHVSKSTKEWMQQKNLLDKIMKTPASSPDINPIENVW
TLEWI-2_CGi     PKHSYKVNVWAGISYKGKTSICIFTGIMDSEIYQKILRDNFVTVTSE--NFPNGYRLYQDNDSKHTSKSTKAWMEDNGILDHIMKTPASSPDLNPIENVW
TLEWI-3_CGi     PKHAYKVNVWAGISYQGKTPICIFTGIMNSVRYQQILESNLLPFVRHRGRFPGGFRLYQDNDSKHTSRSTKAWMEQAGIINSVMKTPASSPDLNPIENVW
TLEWI-4_CGi     PKHAYTVHVWAGISYRERTSICIFNGVMDSVVYQDILDKNCIPFVEE--RFPDGYRLYQDNDSKHKSKSTRDWMAEKGILDNVMTTPASSPDLNPIENVW
TLEWI-5_CGi     PKHSYKVNVWGGISHRGRTSICIFSGIMDSEIYQKILEDNLLPFATA--NYPDGFRLYQDNDRKHTSKSTRRWMEEKQLTESVMTTPASSPDLNPIENVW
TLEWI-1_CVi     PKHAYKVHVWAGISYRGRTPICIFNGIMDSLIYQNILEKNFLPFVED--KFPDGYRLYQDNDSKHKSKSTQDWMAEKGMLQNVMTTPASSPDLNPIENVW
TLEWI-2_CVi     PKHAYKVHVWGGISYEGRTDICIFTGTMDSVTYQHILEKNLIP-ANK--KFPNGFRLYQDNDSKHVSKSTKEWMQQRNLLDKVMKTPASSPDINPIENVW
TLEWI-1_MGall   PKHSYKVHVWGGISYRGRTDVCIFTGIMDSVIYQQILASNFVPFVSA--AFPDGYRLYQDNDSKHKSKSTKEWMRQHGVLDSVMETPASSPDLNPIENVW
TLEWI-1_MPh     PKHAYKVHVWGGISYRGKTDVCCFTGVMDSITYRGILKNNFLPFVNE--RFPDGYRLYQDNDSKHKSKSTVAWVTENGVIDHVMECPASSPDINAIENLW
TLEWI-2_MPh     PKHSYKVHVWGGISYRGRTDICVFTGIMDSIIYQGILEDCLLGFTGS--VFPDGYRLYQDNDSKHKSKSTQEWMRKENMLDKVMEVPASSPDLNPIENVW
TLEWI-3_MPh     PKHSYKVHVWGGVSYRGRTNICVFNGIMDSIIYQQILENNFLPFVNR--AFPDGYRLYQDNDSKHTSKSTKAWMAQNGILDNVMETPASSPDLNPIENVW
TLEWI-1_MYe     PKHAYKVHVWGGISYRGRTSICIFSGIMDSVIYQGILEKKLIPFAER--AFPEGYRLYQDNDSKHKSASTRSWMENHGVLDHVLETPASSPDINCIENVW
TLEWI-4_MYe     PKHAYKV------SYRGRTDICIFSGIMDSIIYKGILQDNLLGFVDA--NFRAGFRLYQDNDRKHTSRSTQDWMRKKGILDSVMSVPASSPDLNPIENVW
TLEWI-1_PMa     PKHAYKVNVWAGISYRGRTSICIFHGIMDSLTYQEILKDNLLPFVEK--KFPEGFRLYQDNDSKHNSRSTKAWYEDNNLTDSIMSTAASSPDLNPIENVW
TLEWI-3_PMa     PKHSYKVNVWAGISYRGRTPICIFTGIMDSVIFQQILQDNLLPFVQR--EFIDGFRLYQDNDSKHNSRSTQEWMRQVGILDTVMKTPASSPDLNPIENVW
TLEWI-4_PMa     PKHAYKVHVWGGISYHGRTDICIFTGTMDSVIYQQILEKNLIPFAKR--VFPNGYRLYQDNDSKHKSKSTKEWMSRNGIEKNFLEAPASSPDINPIENLW
TLEWI-1_PMax    PKHAYKVHVWGGISYSGRTDICIFSGTMDSVIYQSILQDNLLKFTAT--NFGDDFRLYQDNDRKHTSKSTQEWMRQNGILDKVMVVPASSPDLNPIENVW
                          ~87(81-89)                                                     36
                   *       **     *   *  **  *    *   * * * * *
TLEWI-1_BPl     SALKGHLLKEVKPKTKDELIGGIRTFWESLTKEKCCSYIDHIHRVIPFV-----------ILNGGGPSDF------------------------
TLEWI-1_CGi     AAMKLYLTKTVKPKKKEELVAGIRDFWNGLSADGCQKFIDHIHKVIPAV-----------VLNSGEQSGY------------------------
TLEWI-2_CGi     ASMKQYLSNVHKPRTRDELVGGIKAFWASLTAAQCGRYIDHIHRVIPYI-----------VLNKGDASGF------------------------
TLEWI-3_CGi     AGMKTFLQSTAKPKTKEELVSGIRSFWRTLTARQCARYIDHVHRVIPQIRDYLTSSLSLSMAISRYPPCFPLRHISLASLVAAMAIALVPLDQI
TLEWI-4_CGi     ASMKYYLQTVAKPRTKDQLIQGIKAFWETLSTPQCERYIDHIHKVIPYV-----------VLNRGEASGY------------------------
TLEWI-5_CGi     SAMKTYLRSVVKPKKKEELVQGIVSFWNTLSAEKCGHYIDHVHRVIPHV-----------VLNAGGPTQF------------------------
TLEWI-1_CVi     ASMKYYLQTVAKPRTKDQLIQRIRDFWGSLSTAQCGRYIDHIHKVIPVI-----------VLNDGEASGF------------------------
TLEWI-2_CVi     SAMKGFLSRTVKPRKRDELVDGIKRFWEGLSAQSCQRYIDHIHRVIPIV-----------ILNDGEASGF------------------------
TLEWI-1_MGall   AAMKQHLQNKVKPRKKDELVQGIADFWLILTPQKCSAYIDHLHRVLPYV-----------ILNHGGPSGF------------------------
TLEWI-1_MPh     STLKRHLGNVVKPKKKDELVNGIKEFWESLTPEDCSNYIDHIHRVLPTI-----------VLNDGAPSGF------------------------
TLEWI-2_MPh     AAMKQHLQNVVKPKKKDELVKGILDFWSSLSSDNCGSYIDHLHNVIPTV-----------VINEGGPSGY------------------------
TLEWI-3_MPh     AALKMHLENKVKPKKKDELVQGIVDFWTTLTPERCAPYIDHIHRVLPAV-----------VLNHGHPTKF------------------------
TLEWI-1_MYe     ASLKYHLQTVVKPKKKDELVCGIRDFWTSLSTEQCRRYIDHIHKVIPAV-----------ILSDGGPTGF------------------------
TLEWI-4_MYe     SAMKDFIQRQVKPKKKDELIKGIVQFWNDLTPEKCRRFIDHIHRVIPHV-----------VLNRGGPTQF------------------------
TLEWI-1_PMa     ASMKLYLQTVHKPKTRDELVSGIRAFWESLTPEKCAKYIDHIHKVIPYV-----------VLNDGEASGF------------------------
TLEWI-3_PMa     ASLKLHLQTKVKPKRKEDLVRGIQEFWKSLTPESCRKYIEHIHKVIPHV-----------ILNNGDPTDF------------------------
TLEWI-4_PMa     AALKRFLSCEVKPKKKDELVEGIRTFWESLTVDTCSSYIDHIHKVLPHV-----------VLNNGHATKY------------------------
TLEWI-1_PMax    SAMKDQIQRHVKPKKKNELISGIIDFWKGLTPEKCQRFIDHVHRVIPHV-----------VLNRGGPTQF------------------------
```

**Online Resource 3.** Multiple alignment of *TLEWI* transposase sequences. Three α-helices of PAI sub-domain are typed in pink and three α-helices of RED sub-domain are typed in blue. The putative NLS is shaded in gray. The DDE triad of the catalytic domain is shaded in green, and the G-rich box of the DDE domain is shaded in yellow
